# Probing the content of semantic representations in body-selective regions

**DOI:** 10.1101/2025.08.19.669677

**Authors:** Ryuto Yashiro, Masataka Sawayama, Ayumu Yamashita, Kaoru Amano

## Abstract

The extrastriate body area (EBA) has been known to selectively respond to human body parts, but recent work suggests it may encode richer scene semantics. However, it remains unclear what aspects of natural scenes constitute the semantic representation in the EBA. Here we address this question by analyzing the relationship between object co-occurrence in natural scene captions and EBA responses predicted by caption-based encoding models. This revealed object category pairs associated with EBA activity, leading to the hypothesis that EBA encodes the speed of human body motion implied in scenes. A subsequent behavioral experiment and correlation analyses confirmed this, showing strong correlations between EBA responses and implied motion ratings. In addition, the representation of implied body motion extends into the fusiform body area (FBA). Variance partitioning further showed that while body-related features such as the number of people and body size also contributed, implied motion uniquely explained the largest portion of EBA and FBA responses. Overall, semantic representations in body-selective regions are jointly shaped by multiple body-related features, with implied motion as the primary contributor. Our co-occurrence-based approach, combined with brain-to-model mapping methods, offers a novel, interpretable framework for understanding high-level visual representations in natural scene perception.

## Introduction

Recognizing other individuals is fundamental to human social interaction in highly social environments. Among many visual cues, body parts play a crucial role in conveying information regarding individuals’ identities, actions, intentions, and even emotions. Through hierarchical processing over multiple brain regions, the visual system has a remarkable capacity to recognize human bodies accurately^1,2^. This capacity is primarily underpinned by the extrastriate body area (EBA), which has a stronger response to human body parts than to faces and objects^3^. fMRI studies using simplified and well-controlled body stimuli, such as static images of segmented body parts^3–5^ and point-light biological motion^3,6–8^, have demonstrated that EBA selectively processes a form of the human body.

In addition to this conventional approach, neuroscientists have increasingly examined brain responses to more naturalistic visual stimuli, such as images and movies that contain multiple people and objects^9,10^. This approach better approximates the complexity and richness of body recognition in the real world, as human bodies rarely occur in isolation but rather accompany other diverse objects. Recent developments in large-scale fMRI datasets^11–13^ and machine learning models^14–16^ have further facilitated this naturalistic approach, leading to the discovery of neural representations that have been overlooked with the conventional approach using a limited number of simplified stimuli^17–20^. For example, recent studies utilized large language models (LLMs), which generate high-dimensional feature vectors (embeddings) from the text description (caption) of natural images as input, and modeled the brain response to those images based on the corresponding embeddings^21–24^. Importantly, this caption-based encoding model showed a greater capacity to predict brain responses across a wide range of high-level visual regions, including EBA, than encoding models based on visual features extracted from convolutional neural networks^22,24^, or categorical encoding models based on embeddings that capture only the presence of a single object category in an image^24–26^. These findings demonstrate that EBA represents not only a static form of human bodies but also complex semantic contents of an overall scene that the corresponding caption can capture. More specifically, natural scene images containing human bodies do not always elicit a maximum response in EBA; rather, the EBA response varies depending on the semantic context in which the bodies exist.

Despite this new perspective of the neural representation in EBA beyond the traditional view of body selectivity, it remains unclear what information constitutes the semantic representation in EBA owing to the lack of interpretability of caption embeddings. To gain a deeper understanding of the semantic representation in EBA, we consider that the co-occurrence of multiple objects in a scene generally shapes the semantic content of that scene. This motivated the development of a novel analysis aimed at linking the co-occurrence statistics of natural scene captions with corresponding EBA responses.

Specifically, we used large-scale fMRI responses to natural images in the Natural Scenes Dataset (NSD)^12^ along with MS-COCO captions that describe those images^27,28^, and constructed a matrix that contains the number of co-occurrences for multiple pairs of object categories in a group of captions (hereinafter referred to as the co-occurrence matrix; Figure 1A). We then applied matrix factorization to explore which co-occurring categories were key determinants of EBA responses. This data-driven analysis revealed that EBA responses are modulated by the type of object category that co-occurs with human bodies in a natural scene. Qualitative assessment of images with frequently co-occurring categories suggested that dynamic human actions—implied by these objects— may influence EBA responses. To test this possibility rigorously, we conducted a behavioral experiment to obtain subjective ratings of the speed of body motion implied in natural scene images. We computed the correlation between the ratings and brain responses. We found that responses in a large portion of the EBA were highly correlated with implied human body motion and that a region in the fusiform gyrus, commonly known as the fusiform body area (FBA)^29^, showed a similar trend.

**Figure 1.**
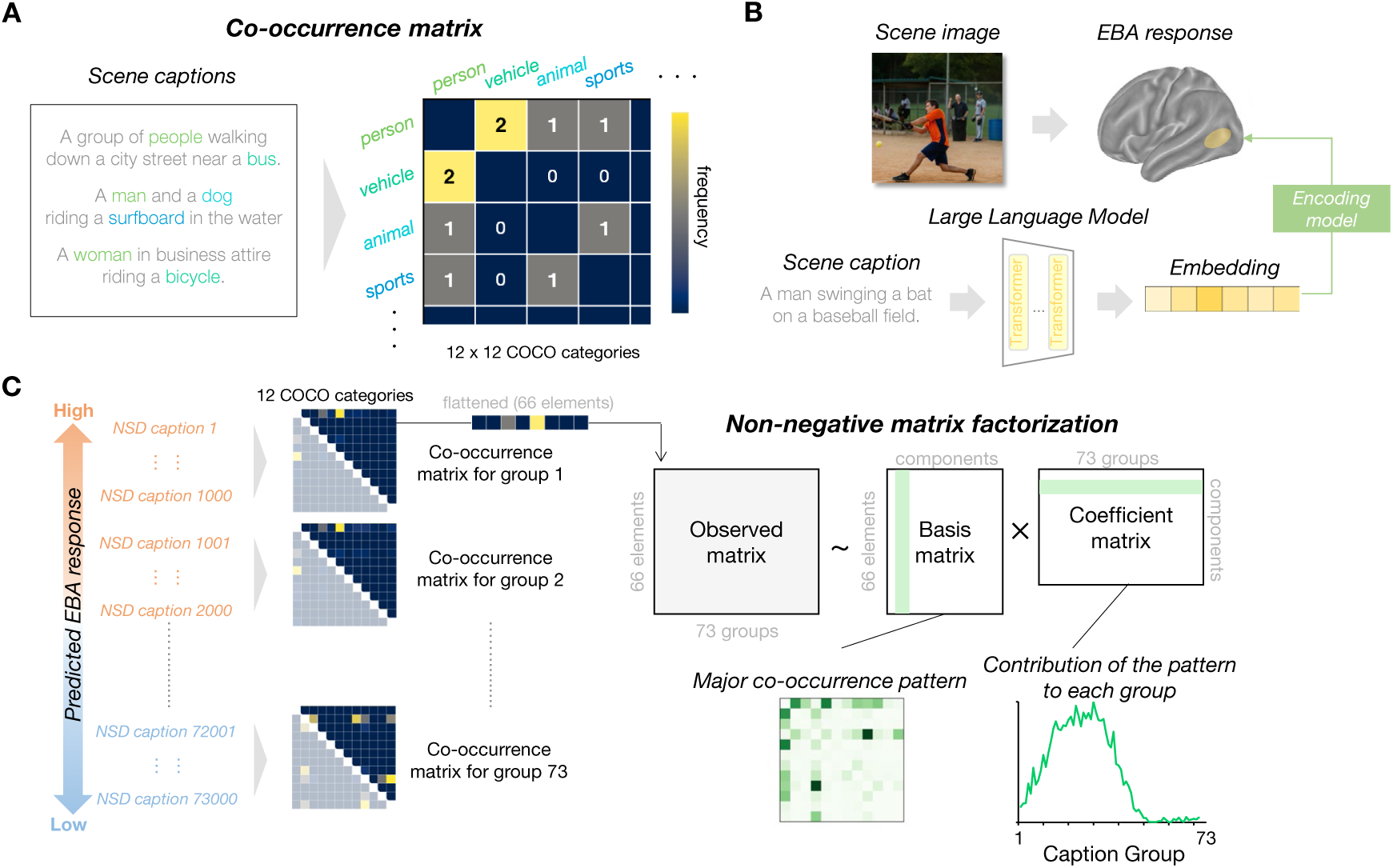
Illustration of our co-occurrence analysis comprising three key elements. (A) Co-occurrence matrix. In our framework, a co-occurrence matrix was defined by counting the co-occurrence frequency of all pairs of 12 MS-COCO superordinate categories in a set of captions that describe natural scene images. By exploring the relationship between the co-occurrence matrix derived from scene captions and EBA responses to those scenes, we sought to gain insights into what specific aspect of semantic content of natural scenes is associated with the representation in EBA. (B) Caption-based encoding model. We constructed encoding models (ridge regression models) using 9000–10000 pairs of image-derived captions and image-evoked responses for each of the eight NSD subjects. Captions were passed through a large language model (MPNet) into 768-dimensional vectors (embeddings), and these embeddings were used as predictors in our encoding models. For copyright reasons, the example scene image is a generated image that preserves the semantic content and composition of the original COCO image. (C) Non-negative matrix factorization. After training the eight encoding models, predicted EBA responses to 73,000 images were calculated for each subject and then averaged across subjects. Based on these averaged predicted responses, all captions were sorted and divided into 73 groups. For each group, a co-occurrence matrix was constructed. The 66 upper-triangular elements from each matrix were flattened and concatenated column-wise to form a single 66 × 73 matrix. We then performed non-negative matrix factorization to decompose the observed matrix into two matrices: basis and coefficient matrix. Each column of the basis matrix represents co-occurrence patterns for each component when reshaped back into its original two-dimensional form, while each row of the coefficient matrix (i.e. a coefficient vector) reflects how strongly each co-occurrence pattern is represented across the 73 groups. Since the groups were sorted in descending order of predicted EBA response, each coefficient vector captures the relationship between the corresponding co-occurrence pattern and the magnitude of predicted EBA responses. The optimal number of components was determined by minimizing the Bayesian Information Criterion (BIC).

Furthermore, we computed other body-related features for each image and performed variance partitioning to disentangle the unique and shared contributions of each feature to the representation in EBA and FBA. The results showed that responses in substantial portions of body-selective regions were uniquely explained by implied motion, whereas other features, such as the number of people and body size in an image, also uniquely accounted for the response in small but spatially distinct clusters within these regions.

Overall, our findings suggest that multiple body-related features jointly constitute the semantic representation in EBA and FBA, with implied motion serving as the primary contributor. We argue that our framework, which combines co-occurrence analysis and existing brain-to-model mapping methods such as encoding models and variance partitioning, provides a promising approach for better interpreting high-level neural representations in the brain.

## Results

### Data-driven analysis of object co-occurrence revealed that object pairs associated with the EBA response

The present study aims to understand precisely what semantic aspects of natural scenes form the neural representation in EBA. To this end, we first focused on the fact that the joint presence of multiple objects generally shapes the overall semantic content of a scene. For example, a person sitting at a table with a fork and a dish may be interpreted as preparing to eat, whereas replacing those items with a laptop leads us to interpret the scene as someone working. Building on this idea, we developed a novel analysis of the relationships between the co-occurrence patterns of objects in large-scale natural scene images and the corresponding EBA responses (Figure 1). Through this analysis, we sought to reveal which pairs of object categories are associated with the magnitude of the EBA response and to characterize the semantic information represented in EBA in an interpretable manner.

Our co-occurrence analysis framework is centered around a matrix called a co-occurrence matrix. Each element in this matrix reflects the co-occurrence frequency of words of multiple object categories in a set of captions that provide textual descriptions of natural scenes (see Figure 1A as an example). We used 12 MS COCO superordinate object categories: accessory, animal, appliance, electronic, food, furniture, indoor, kitchen, outdoor, person, sports, and vehicle (see Supplementary Table 1 for subordinate categories for each of the 12 categories). As a whole, the matrix captures important information regarding the co-occurrence patterns that are associated with brain responses to those scenes. One rationale for using image captions instead of the objects depicted in the images themselves is that images often contain too many objects, some of which may be irrelevant to the overall semantic content of a scene (e.g., a small cup on a desk when people have a meeting in a room). In contrast, captions capture the essential components of a scene, providing information that aligns closely with human semantic scene recognition.

Our co-occurrence analysis involved the following steps. First, we trained an encoding model using large-scale fMRI responses to natural images for each of the eight human subjects in the Natural Scenes Dataset (NSD)^12^, along with corresponding human-annotated captions of the scene images (Figure 1B). We obtained caption embeddings from a large language model^30^ and used them to fit ridge regression models that predict EBA responses averaged across vertices for each subject. Next, we used these trained models to predict EBA responses to all 73,000 captions, sorted these captions in descending order of the average predicted EBA responses across the eight subjects, and divided these captions into 73 groups of 1,000 captions. For each group, we constructed a co-occurrence matrix that consists of the number of pairwise object category co-occurrences across the captions (Figure 1C). We subsequently perform nonnegative matrix factorization to extract major co-occurrence patterns and their contributions to the original matrix of each caption group. These procedures, which are based on caption-based encoding models and co-occurrence matrices, make full use of the entire set of 73,000 caption`‒`response pairs in the NSD, which far exceeds the 1,000 image`-`evoked responses shared across eight subjects. This enabled the reliable extraction of general object co-occurrence patterns that are systematically associated with EBA responses.

The caption-based encoding models showed a high capacity to predict the overall EBA response for all the subjects (Pearson correlation ranging from 0.44 to 0.72; see Supplementary Table 2 for individual data). Based on the predicted responses from these high-performing encoding models, the co-occurrence matrices for 73 groups of captions were decomposed into three components (hereinafter, components 1 to 3), each associated with high, moderate, and low EBA responses, as shown in the distinct peak location of the corresponding coefficient vector (Figure 2A). The bar plots in Figure 2A show the top 10 word category pairs ranked by co-occurrence frequency for each component. As expected, components 1 and 2 (associated with high and moderate EBA response) included the ‘person’ category, whereas it was absent from most category pairs in component 3 (related to low EBA response). We also found that the ‘animal’ category frequently occurred in component 2 (moderate EBA response). This finding is consistent with previous findings that EBA shows weaker responses to animal bodies than to human^3^, supporting the validity of our method for revealing the functional properties of the body-selective regions.

**Figure 2.**
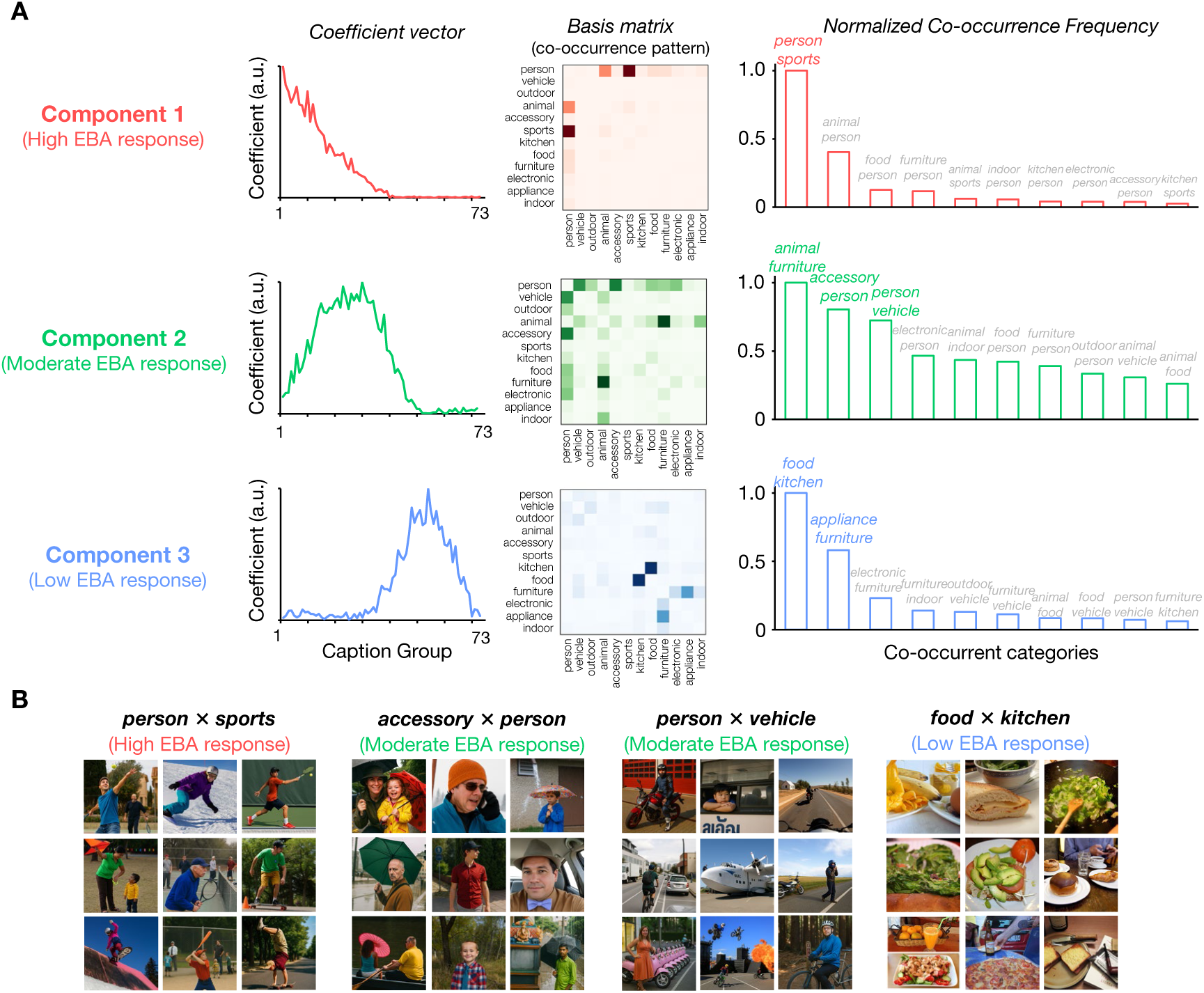
Results of the co-occurrence analysis. (A) Three major co-occurrence components were revealed by non-negative matrix factorization applied to the 73 co-occurrence matrices. Each component is associated with high, moderate, and low EBA responses, respectively, as indicated by the different peak locations of the coefficient vectors along the horizontal axis (left), which represents the caption groups sorted by predicted EBA response magnitudes. The middle panels show the co-occurrence pattern for each component, which are summarized as the bar plots of normalized co-occurrence frequency of the top-10 co-occurring category pairs (right). The pair labels are colored if the corresponding value exceeds 0.5. The category ‘person’ co-occurred with ‘sports’ in component 1, and with ‘accessory’ and ‘vehicle’ in component 2, suggesting that EBA response varies depending on the object categories co-occurring with human bodies. (B) Example COCO images containing the major co-occurring category pairs. Qualitative inspection of these images indicates that EBA tends to respond strongly to images implying dynamic human body motion, moderately to those containing static human bodies, and weakly to those without human bodies. For copyright reasons, COCO images containing human faces have been replaced with images generated by ChatGPT-5, which preserve the semantic content and composition of the original images.

Notably, the ‘person’ category co-occurred with distinct categories across components: with ‘sports’ in component 1 and with ‘accessory’ and ‘vehicle’ in component 2. We then qualitatively assessed images whose captions contained these co-occurring categories (Figure 2B). We found that images associated with higher EBA responses tend to imply more dynamic human body motion (e.g., hitting a ball with a racket or doing acrobatics). In contrast, those associated with moderate EBA responses tend to contain a static form of the human body (e.g., standing still with an umbrella). These results raise the possibility that EBA encodes human body motion implied by co-occurring objects in a natural scene.

### Behavioral experiment collecting subjective ratings of implied body motion

To test the possibility that EBA represents implied human body motion in a natural scene, we next sought to assess the correlation between the magnitude of EBA responses and the speed of implied motion^31,32^ across NSD images. As the NSD does not provide information regarding the speed of implied body motion for each image, we conducted a behavioral experiment for five human participants to collect their subjective ratings of implied motion speed for the NSD images, which were used in the subsequent correlation analyses.

We subsampled 100 NSD images containing more than one human body, excluding those that contained both a person and an animal or both a person and a vehicle, as these could imply motion not solely attributable to the human body (for example, we excluded images of a person riding a horse). To make participants aware of the range of implied body motion speed across images, the experiment started with a passive-viewing block, in which participants observed all 100 person images without making any response, each successively presented for 1 sec. In subsequent rating blocks, each image was presented for 3 s, during which participants rated the implied motion of the person depicted in the image on a scale from 1 to 5. We explicitly instructed participants on the rating criteria: 1 indicates static motion, and 5 corresponds to the fastest motion they had previously seen in the passive-viewing session. Each image was presented twice throughout the experiment to assess the within-participant reliability of the ratings.

All five participants showed consistency in the ratings, as indicated by high within-participant reliability across two repetitions (Spearman rank correlation ranging from 0.85 to 0.96; Supplementary Figure 1). The between-participant reliability was also high across all participant pairs (Spearman rank correlation ranging from 0.80 to 0.90, Supplementary Figure 2), suggesting that the participants provided highly similar ratings. These high reliability scores justified averaging the ratings across participants as an estimate of the general population, which can be compared to the fMRI responses obtained from a different participant group in the NSD. We thus averaged the ratings across participants and repetitions to obtain one representative rating of the implied body motion for each image (Figure 3A). The participants rated the implied body motion in the images on the basis not only of their category but also of the broad semantic context in which the person appeared. Indeed, images rated as implying high motion speed frequently included a person with sports equipment, yet some sports-related images received low ratings when the person in those images did not substantially move or merely stood still.

**Figure 3.**
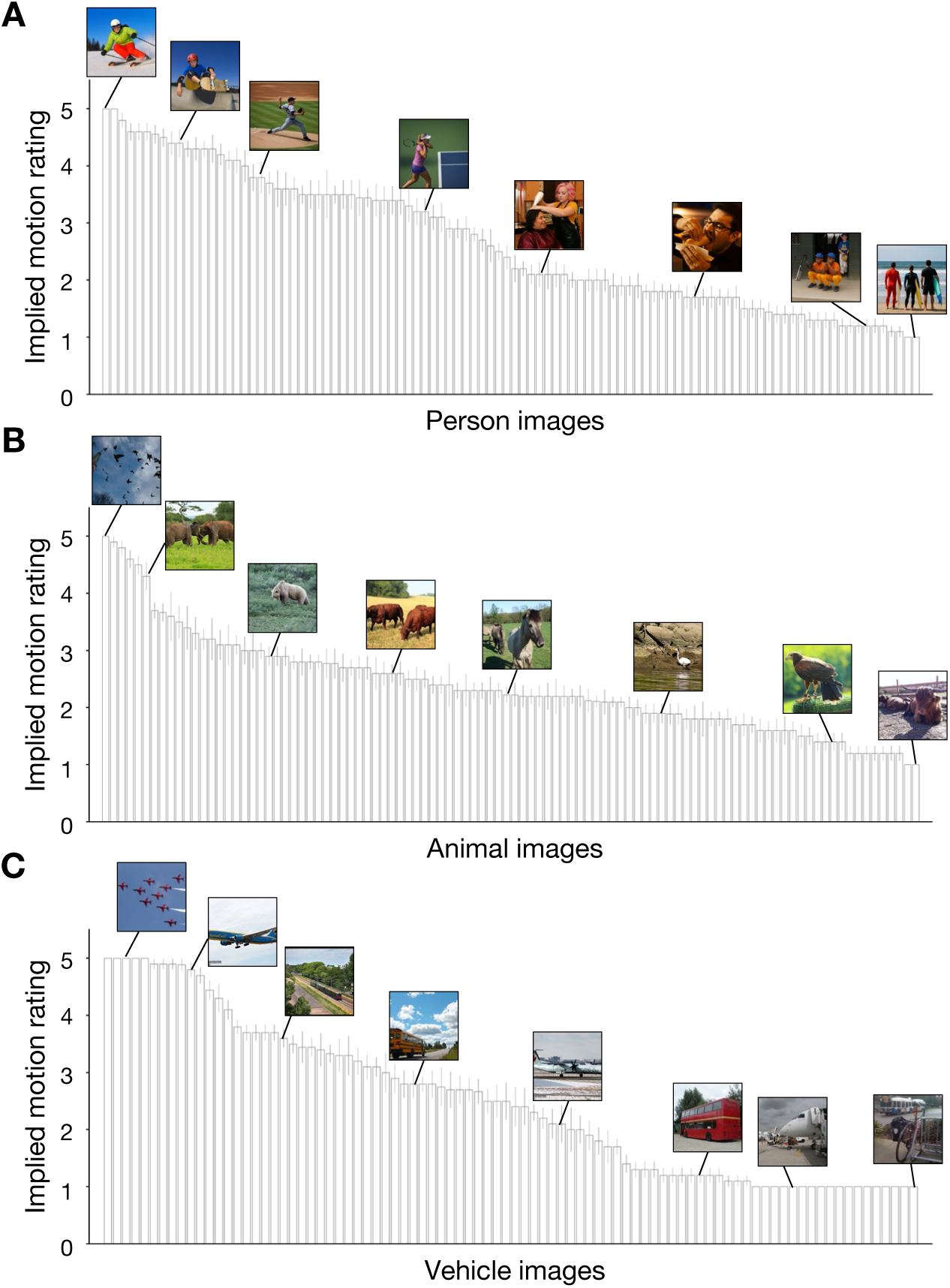
Subjective ratings of implied motion from our behavioral experiment. Bar plots show ratings averaged across five participants and two repetitions for person (A), animal (B), and vehicle (C) depicted in natural scene images. Ratings were broadly distributed across the full range, from 1 (static) to 5 (fastest motion of all images). Images within the same object category (e.g., sports-related scenes, birds, and airplanes) were rated as implying different levels of motion speed, indicating that participants relied on contextual information rather than category membership alone. For copyright reasons, COCO images containing human faces have been replaced with images generated by ChatGPT-5, which preserve the semantic content and composition of the original images.

Additionally, we investigated whether EBA also represents the implied motion of animal bodies or even objects. To test this possibility, we conducted the same rating experiment using multiple NSD images of animals and vehicles to collect implied motion ratings for these categories. Again, we carefully selected animal and vehicle images, ensuring that all the animal (vehicle) images did not contain people or vehicles (animals). The within-participant reliability for these two types of images was comparable to that for the person images (Spearman rank correlation ranging from 0.67 to 0.93 for the animal images and 0.89 to 0.96 for the vehicle images; Supplementary Figure 1). The between-participant reliability for the animal images was lower than that for the person and vehicle images, but was still significantly high (Spearman rank correlation ranging from 0.40 to 0.71 for the animal images and 0.81 to 0.89 for the vehicle images; Supplementary Figure 2). Thus, we again obtained one average rating for each image and found that the ratings were widely distributed across the animal and vehicle images (Figure 3B and 3C).

### Responses in body-selective regions are highly correlated with implied motion ratings of natural scene images

Having collected the implied motion ratings for people, animals, and vehicles in natural scene images, we used these ratings to investigate the neural correlates of implied motion, with a primary focus on EBA. After defining the EBA for each NSD subject via cortical surface t-statistic maps derived from functional localizer experiments in the NSD, we examined whether higher EBA responses were associated with higher implied motion ratings for person, animal, and vehicle images. Specifically, we computed overall EBA responses averaged across vertices and performed linear mixed-effects regression separately for each of the three categories, with implied motion ratings as fixed effects and subjects as random effects. We found that the EBA responses were significantly correlated with the implied motion ratings for person images (beta=0.040, SE=0.012, *p*=0.001, CI=[0.016, 0.063]), which is consistent with our hypothesis from the qualitative observation in the co-occurrence analysis (Figure 2B). For animal images, a significant correlation was also observed between the overall EBA responses and ratings (beta=0.046, SE=0.013, *p* < 0.001, CI=[0.022, 0.071]). However, the implied motion ratings for vehicle images did not explain the overall EBA responses (beta=-0.006, SE=0.008, *p*=0.426, CI=[-0.022, 0.009]).

Is the observed correlation with implied body motion solely attributable to EBA? Previous studies have shown that activation in two motion-processing regions, the middle temporal (MT) and middle superior temporal (MST) regions, is modulated by body motion speed implied in natural scenes^33–35^. As these regions are anatomically proximal to or partially overlap with EBA^6,36,37^, the observed correlation may be fully explained by activity in these motion-processing regions. Indeed, for NSD subjects, some EBA vertices were also labeled MT or MST if we defined those regions via a probabilistic atlas (see yellow, green, and blue contours in Figure 4A). To test this possibility, we restricted our analysis to EBA vertices located outside anatomically defined motion-processing regions. A linear mixed-effects regression model was fitted using the same implied motion ratings and responses averaged across the restricted set of vertices. We found that the response in the EBA outside the MT and MST was still significantly correlated with the implied motion rating for person images (beta=0.033, SE=0.014, *p*=0.022, CI=[0.005, 0.061]) and animal images (beta=0.040, SE=0.014, *p*=0.003, CI=[0.013, 0.068]) but not for vehicle images (beta=-0.010, SE=0.008, *p*=0.225, CI=[-0.025, 0.006]). These results indicate that EBA uniquely encodes human and animal body motion implied in static natural scene images.

**Figure 4.**
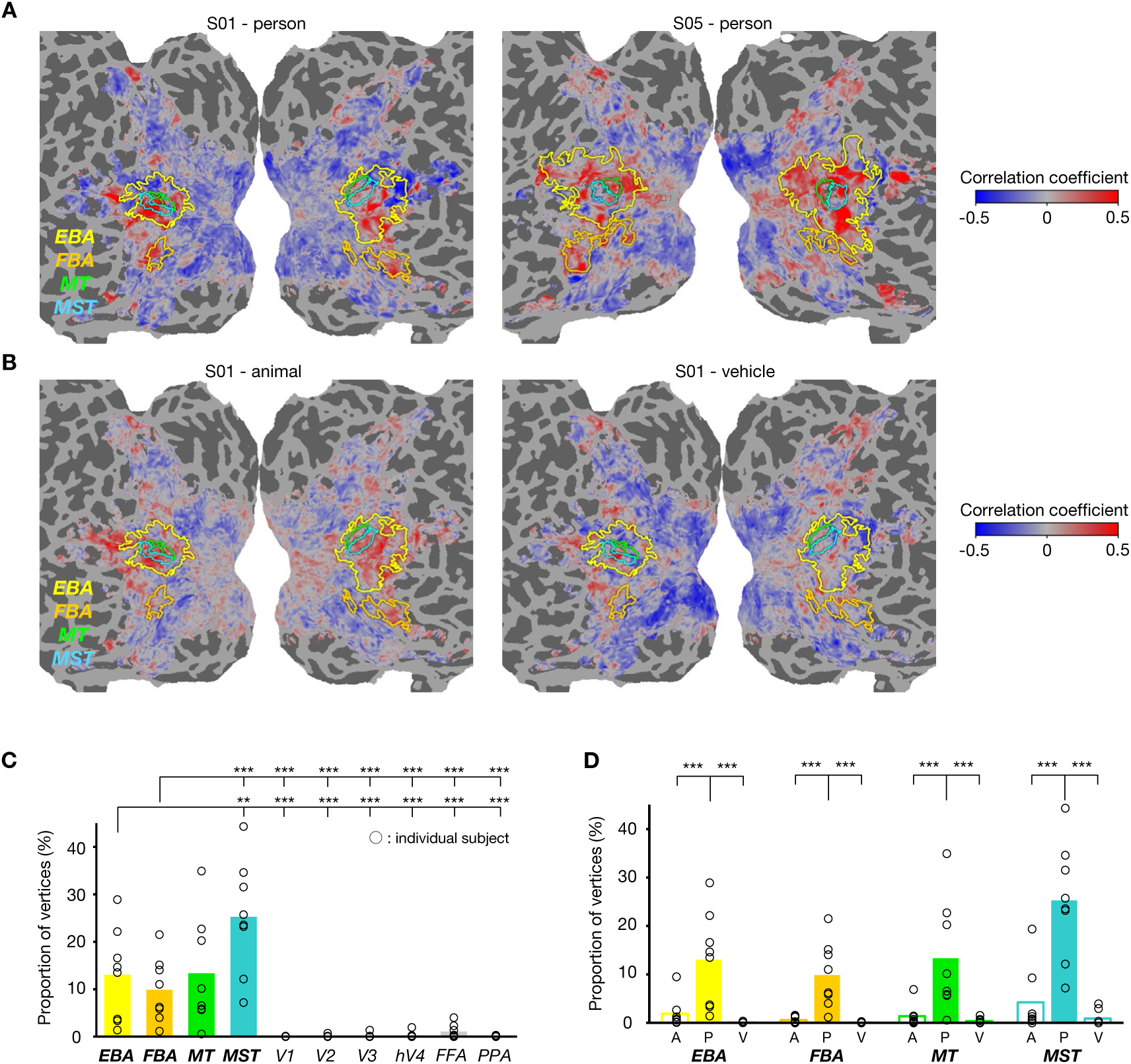
Flattened cortical surface maps of correlation coefficients with implied motion ratings for three image categories (person, animal, and vehicle). (A) Correlation maps for person images from two NSD subjects (subject 01 and 05). Colored contours indicate body-selective (yellow and orange; EBA and FBA) and motion-selective regions (green and blue; MT and MST). A large portion of these regions showed high correlations with the speed of implied human body motion in natural scene images. (B) Correlation maps for the animal and vehicle images from one NSD subject (subject 01). A small portion of vertices in EBA exhibits high correlations with animal implied motion, but not with vehicle implied motion. (C) Proportion of vertices showing significant correlation with implied motion for person images across visual regions. (D) Proportion of vertices showing significant correlation with implied motion for animal (A), person (P), and vehicle (V) images within body-selective and motion-selective regions. The significance of vertex-wise correlations in (C) and (D) was assessed by one-sided permutation tests, with 1000 randomizations of the stimulus labels (FDR corrected *p* < 0.05). Asterisks indicate significant differences in proportions determined by one-sided Wilcoxon signed-rank tests with FDR correction (**: *p* < 0.05, ***: *p* < 0.01).

Next, we aimed to investigate the cortical organization of implied body motion representations within body-selective regions. To this end, we visualized a representational map of implied body motion by computing the correlation between the implied motion ratings for person images and the responses measured at each vertex and mapping the correlation values onto a cortical surface. We computed the correlation across the entire visual cortex, encompassing motion-selective regions (MT and MST), a body-selective region located in the fusiform gyrus (fusiform body area; FBA), and other category-selective regions (fusiform face area^38^ and parahippocampal place area^39^; FFA and PPA), to facilitate comparisons across different visual regions. Figure 4A shows cortical maps of the correlations with implied motion ratings for two representative NSD subjects. We observed that a large portion of the vertices in the EBA, MT, and MST exhibited high correlations.

The vertices with high correlations were broadly distributed across the ventral and dorsal parts of the EBA, with no consistent spatial pattern observed across the eight subjects (correlation maps for all subjects are provided in Supplementary Figure 3). Interestingly, a portion of the FBA, located far from the EBA, MT, and MST, also showed high correlations with implied human body motion. In contrast, regions along the low-level visual cortex and high-level ventral (except for the FBA) and dorsal visual streams presented substantially lower correlation values. These qualitative observations suggest that information regarding implied human body motion is localized in motion-processing and body-selective regions located in the lateral and ventral parts of the brain and is widely distributed within those regions.

To quantitatively compare the spatial extent over which implied body motion is represented across visual regions, we performed a permutation test to compute the proportion of vertices showing significant correlations in each region (Figure 4C). We found that a greater proportion of vertices in the EBA (located outside motion-processing regions) and FBA exhibited significantly greater correlations than those in the low-level and high-level visual regions, including the FFA and PPA (one-tailed Wilcoxon signed-rank tests, *p* < 0.01). The proportion of significantly correlated vertices in the EBA and FBA was comparable to that in the MT (one-tailed Wilcoxon signed-rank tests, *p* > 0.05) but smaller than that in the MST (one-tailed Wilcoxon signed-rank tests, *p* < 0.05). These results suggest that, similar to motion-selective regions, two anatomically distinct body-selective regions—EBA and FBA—represent human body motion implied in static natural scenes.

We next turned to another image category—animals and vehicles—to test whether the body-selective regions also encode the implied motion of these nonhuman categories at the single-vertex level. Figure 4B shows cortical maps of the correlations between the brain responses and implied motion ratings for animals and vehicles obtained from our behavioral experiment (correlation maps for all subjects are provided in Supplementary Figure 4). In contrast to the correlation maps for person images (Figure 4A), these correlation maps, particularly those for vehicle images, did not show a clear distinction between body-selective/motion-selective regions and the surrounding regions. We again assessed the significance of vertex-wise correlations for each category and computed the proportion of vertices with significant correlations within each body-selective and motion-selective region (Figure 4D). The proportion of vertices significantly correlated with the implied motion for the animal and vehicle images was substantially smaller than that for the person images (one-tailed Wilcoxon signed-rank tests, *p* < 0.01). Taken together, these results indicate that EBA and FBA represent the implied motion of both human and animal bodies, but with a greater proportion of vertices showing high correlations for human bodies than for animal bodies.

### Body-selective regions represent other body-related features in addition to implied body motion

Our co-occurrence and correlation analyses suggest that the EBA and FBA play a functional role in representing human body motion, as implied by co-occurring objects in a natural scene. However, these body-selective regions may also encode other aspects of the human body beyond implied motion, as suggested by previous studies. One recent study using the NSD and caption-based encoding models identified functional clusters in EBA that reflect the number of people in an image^40^. Additionally, the response in body-selective regions may be explained simply by the low-level visual properties of the human body in the image. For example, EBA may exhibit different overall response levels depending on the proportion of an image covered by the human body, which is consistent with findings reported in a prior study^4^. It is also possible that the EBA response can be modulated by the position of human bodies relative to the image center, given previous reports that EBA contains information regarding the position of a body stimulus in the visual field^36,41^. To test these possibilities, we extracted additional body-related features from the person images used in our experiment and performed the same correlation analysis using these new features.

We defined three additional body-related features for each person image—body size, number of people, and distance between the image center and the largest body in the image—by using a text-based segmentation model (Grounded Segment Anything)^42^. We provided this model with our person images and the word “person” as input. We obtained output images with segmented areas and bounding boxes (rectangular frames surrounding the segmented bodies) indicating the presence of people in the original images (Figure 5A). For each output image, we obtained one value for each of the three features (see Methods for details). Additionally, the RMS contrast was computed for the original images as a proxy for low-level information regarding spatial frequency. This was motivated by a qualitative observation that images rated as implying higher motion speed tended to contain larger uniform background areas (e.g., blue sky or snow field; Figure 3A). We then computed the correlation between these four features and responses in the body-selective regions and compared the proportion of vertices showing significant correlations across features.

**Figure 5.**
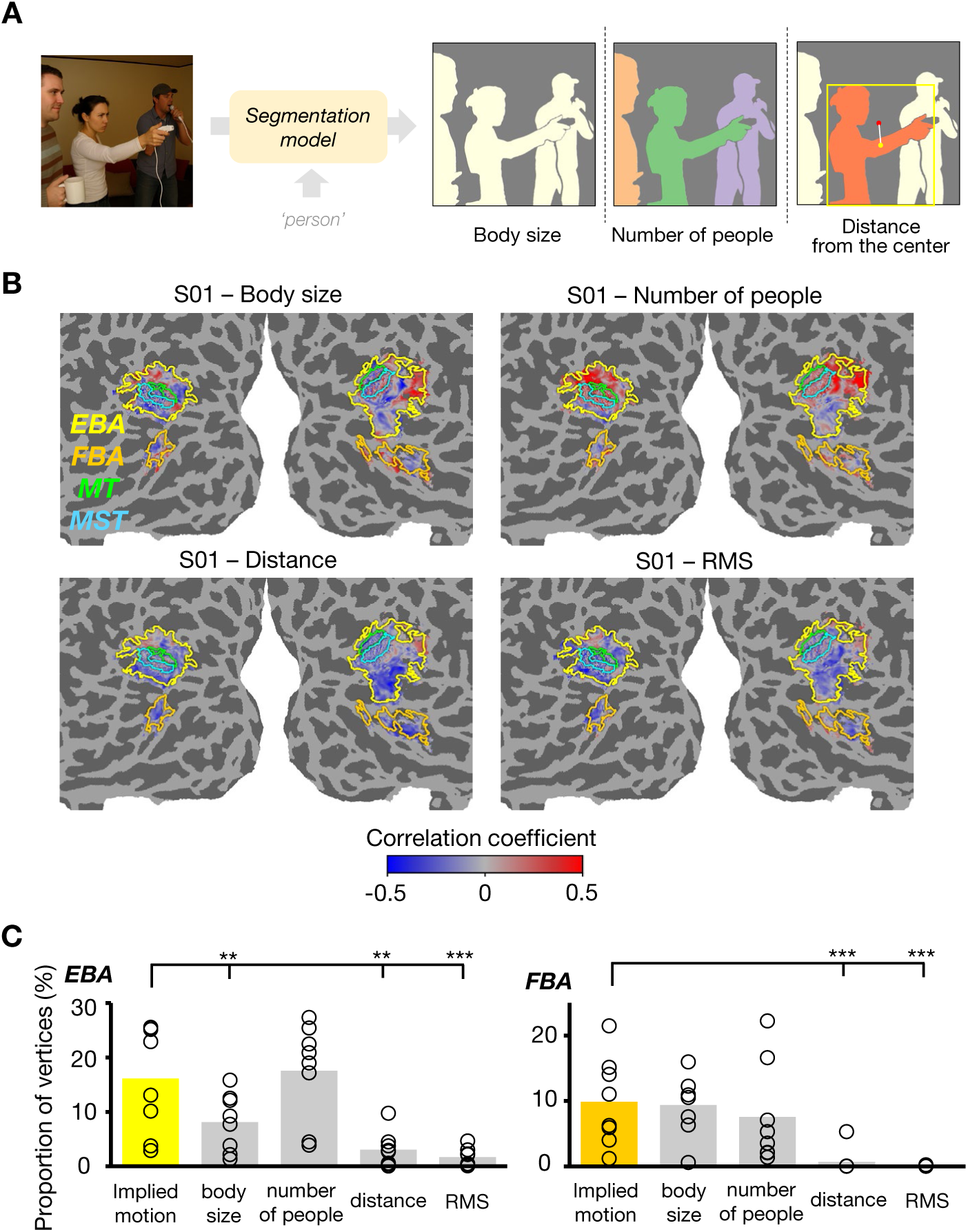
Representation of other body-related features in body-selective regions. (A) A state-of-the-art segmentation model (Grounded Segment Anything) with text input was used to obtain images with segmented human body parts and bounding boxes (a yellow rectangle in the image on the right as an example). Based on these output images, we computed body size as the number of pixels corresponding to bodies, number of people as the number of bounding boxes with ‘person’ labels, and distance from the center as the Euclidean distance between the center of an image and that of the largest bounding box. For copyright reasons, the example image is a generated image that preserves the semantic content and composition of the original COCO image. (B) Flattened cortical surface maps of correlation coefficients with the three body-related features and one low-level visual feature (RMS contrast) for subject 01. Some voxels exhibited high correlations with body size and number of people, whereas distance from the center and RMS contrast explained little of the responses for most vertices. (C) Proportion of vertices in EBA (left panel) and FBA (right panel) showing significant correlations for each feature. The significance of correlations was assessed by two-sided permutation tests with 1000 randomizations of the stimulus labels (FDR corrected *p* < 0.05).

Figure 5B shows flattened surface maps of the correlations with the four additional features for one representative subject (correlation maps for all the subjects are provided in Supplementary Figure 5). Clusters of high correlations were observed for body size and number of people, whereas correlations for distance from the center and RMS contrast were close to zero for most vertices. To quantitatively compare the cluster size of significant correlations across features, we again computed the proportion of vertices significantly correlated with each feature for EBA and FBA (Figure 5C). We found that for both EBA and FBA, approximately 10–15% of vertices on average exhibited significantly high correlations with implied motion, body size, and the number of people. Statistical comparison indicated that implied motion was correlated with EBA and FBA responses in larger proportions of vertices than with distance from the image center and RMS contrast (two-sided Wilcoxon signed-rank tests, *p* < 0.05). We did not observe a significant difference in the proportion of vertices between implied motion and the number of people in the EBA or between implied motion and body size/number of people in the FBA (two-sided Wilcoxon signed-rank tests, *p* > 0.05). These results indicate that other body-related features are also represented in body-selective regions: the number of people in the EBA and FBA, and the body size in the FBA, each to a degree comparable to the representation of implied body motion. Asterisks indicate significant differences in proportions, assessed by two-sided Wilcoxon signed-rank tests with FDR correction (**: *p* < 0.05, ***: *p* < 0.01).

### Variance partitioning identified a larger functional cluster of implied body motion and smaller clusters of other body-related features

Our results suggest that responses in body-selective regions can be captured by multiple body-related features, especially implied body motion, body size, and the number of people in natural scenes. This leads to the following question: where are such features uniquely represented within these regions? As the body-related features are not orthogonal to each other (a correlation matrix of all pairs of features is provided in Supplementary Figure 6), simply constructing cortical correlation maps and computing the proportion of vertices with significant correlations separately for each feature is insufficient to reveal the fine-grained cortical organization of feature representations. A more appropriate approach is to quantify the amount of response variance uniquely explained by each individual feature. Thus, we performed variance partitioning^43,44^ using the three features most strongly associated with EBA and FBA—implied motion, number of people, and body size—and identified cortical locations where each of these features best accounted for responses.

We fitted seven linear regression models separately, each incorporating either a single feature or a combination of two or three features, and computed the 5-fold cross-validated R-squared values for each model to quantify the variance uniquely explained by individual features and the shared variance jointly explained by feature pairs or triplets (see Methods for details). Figure 6 shows flattened surface maps of the unique and shared variance for one representative subject. A large portion of responses in the body-selective regions was uniquely explained by one of the three features, with each feature best accounting for a distinct set of vertices. In contrast, only a small portion of the response variance was explained by a combination of two or three features. Variance maps for all subjects (Supplementary Figure 7) revealed similar trends, despite variability in the spatial distribution of each feature cluster across subjects.

**Figure 6.**
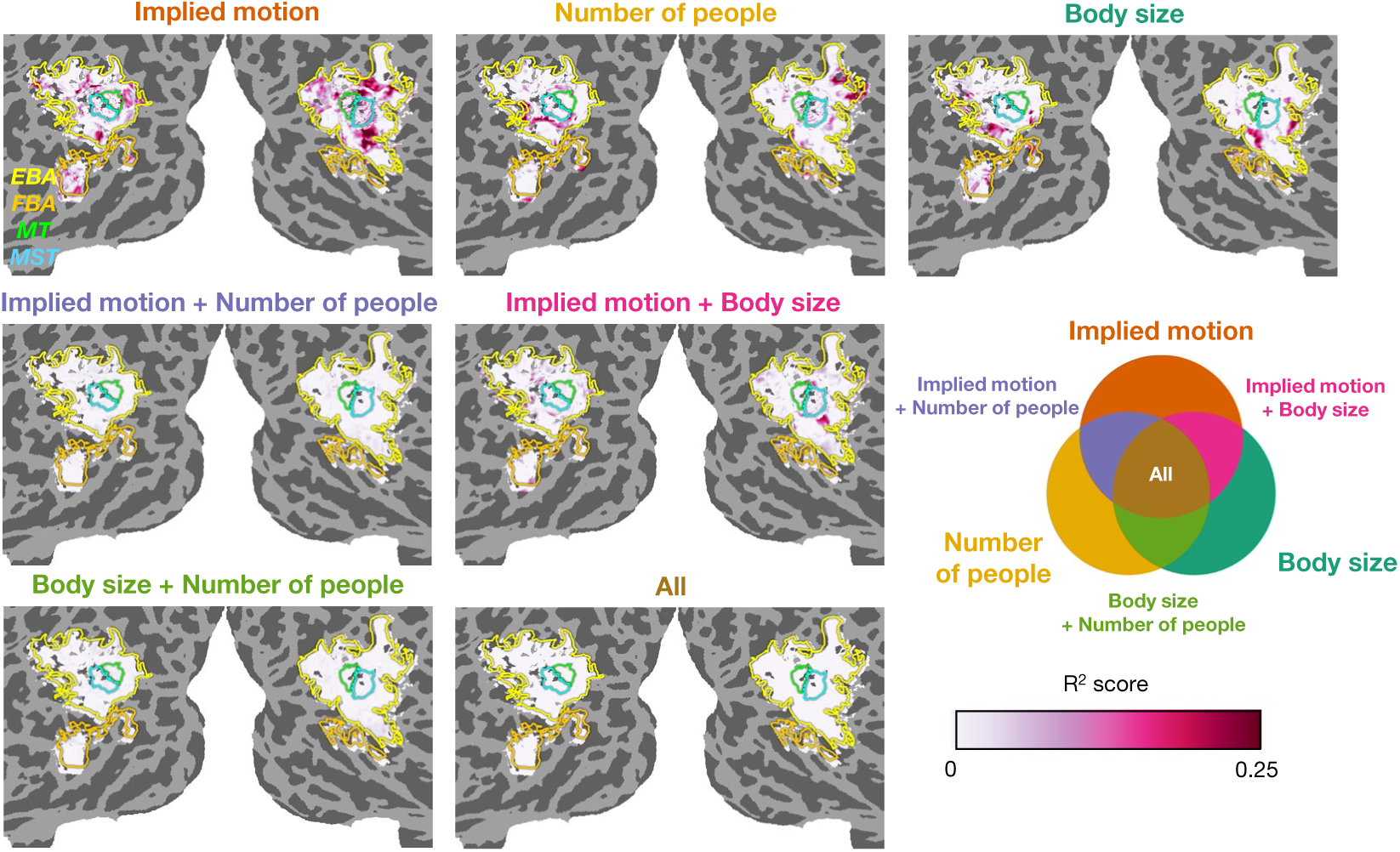
Results of variance partitioning for one subject (subject 05). We computed the variance uniquely explained by each feature, as well as the variance jointly explained by all possible combinations of the three body-related features (implied motion, body size, and number of people). The three panels in the top row show cortical surface maps of R-squared values, which quantify the unique contributions of each individual feature to the responses in EBA and FBA. The bottom four panels show cortical surface maps of R-squared values, which quantify the shared contributions between each pair of features or across all three features. Each feature uniquely explains distinct cortical locations in EBA and FBA, whereas the shared contributions of multiple features are substantially smaller.

To visualize the cortical location of functional clusters that encode each body-related feature, we constructed cortical maps indicating which feature explained the largest fraction of variance for each significantly predicted vertex. We then computed the proportion of vertices best explained by each individual feature or combination of features to quantify the size of each cluster (Figure 7; see Supplementary Figure 8 for all subjects). We found that the majority of vertices were best and uniquely explained by the implied body motion in EBA for 7 out of 8 subjects and in FBA for 5 out of 8 subjects. For some subjects, the second largest proportion of vertices in EBA was best explained by the number of people (subjects 2, 4, 5, and 7). In contrast, body size best accounted for the second largest proportion in FBA (subjects 1, 2, 3, 5, and 7). These results suggest that multiple aspects of human bodies in natural scenes jointly shape the representation in body-selective regions in a complementary manner, with implied body motion serving as the primary contributor.

**Figure 7.**
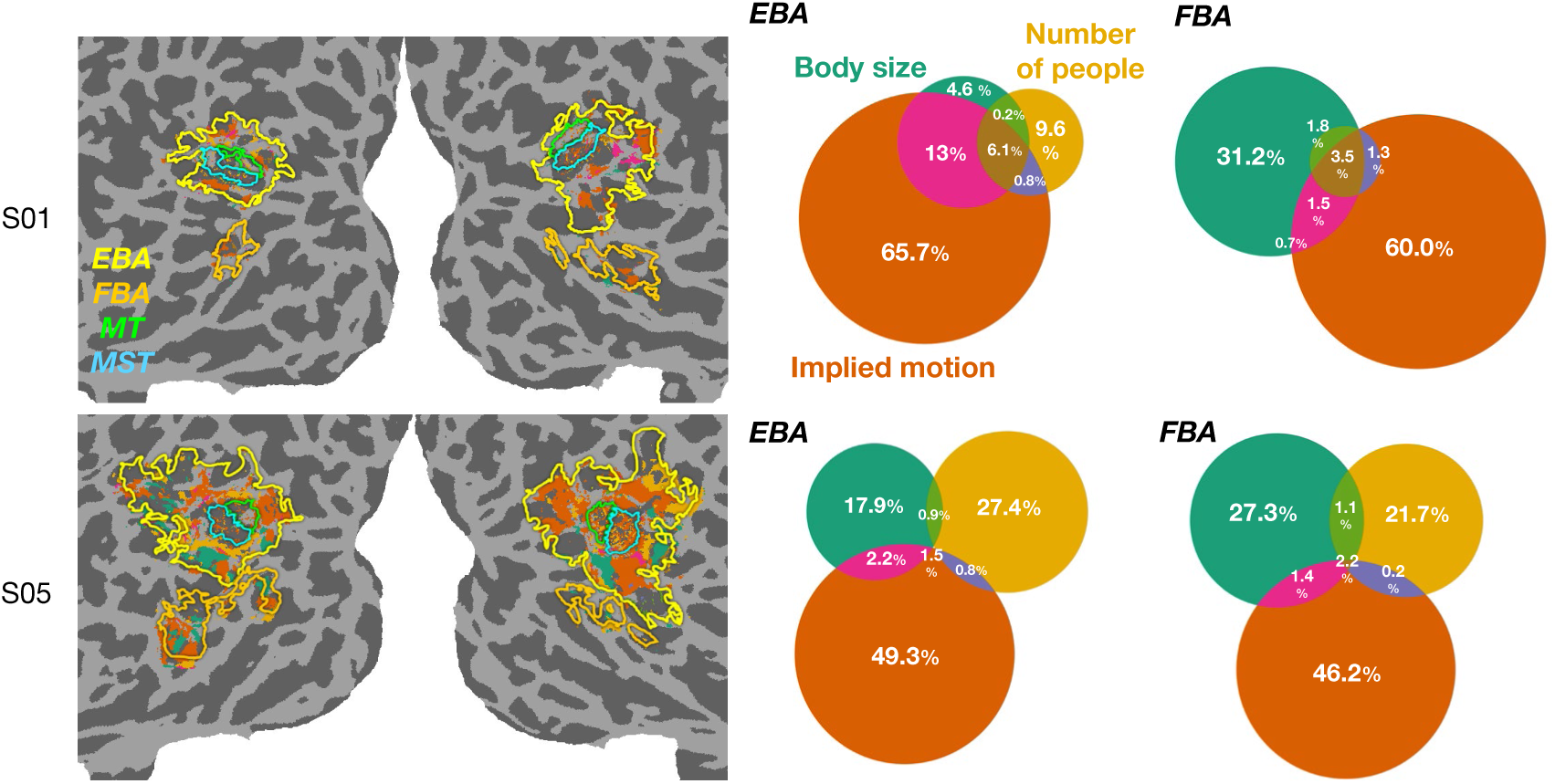
Cortical surface maps of largest variance partition for two subjects. Each significantly predicted vertex is assigned a color corresponding to the body-related feature that explains the largest fraction of variance. The Venn diagrams summarize the proportion of significantly predicted vertices best explained by each feature (note that the size of each area is not scaled to reflect exact proportions). Implied motion uniquely and best explained the majority of vertices in both EBA and FBA.

## Discussion

We investigated semantic representations of body-selective regions by combining several analytical and experimental methods. We first developed a novel analysis focusing on the relationship between the co-occurrence of multiple objects in a wide array of natural scene images and the corresponding brain responses predicted from the caption embeddings of those images. This analysis revealed several category pairs that are associated with the magnitude of the EBA response and suggested that EBA may encode human body motion implied in static scene images. Subsequent correlation analyses corroborated this hypothesis: a large portion of EBA and FBA vertices indeed exhibited significant correlations with subjective ratings of the speed of implied human body motion that were collected in our behavioral experiment. We also derived additional body-related features from person images and disentangled the contribution of each individual feature to the semantic representation in EBA and FBA through variance partitioning. We found that implied motion, number of people, and body size depicted in natural scene images uniquely explain distinct cortical locations of the EBA and FBA, with the largest cluster representing implied human body motion.

One conventional approach to understanding the functional properties of body-selective regions involves measuring brain responses to a few isolated body part stimuli that are hand-picked by experimenters under the assumption that these regions are particularly tuned to a single categorical aspect of the human body. Although this approach has revealed several functional roles of the body-selective regions^4–6,45–48^, the stimuli used in this approach clearly lack diversity and ecological validity, thereby limiting generalizability beyond well-controlled experimental settings^9^. Our study extends this previous approach in two ways. First, we used a large-scale neural dataset (NSD)^12^, which contains high-quality fMRI responses to a large number of natural scene images. This dataset supports the discovery of neural representations that generalize to varied visual input, as demonstrated in several recent studies^18,19^. Second, and more importantly, building on recent findings that category-selective regions represent not only a single object category but also complex semantic contents of a natural scene^22,24^, we focused on the co-occurrence of object categories as a crucial factor shaping the semantic content of a scene, and explored its relationship with responses in EBA. This novel approach enables us to go beyond the traditional view of body selectivity yet still examine in an interpretable way what constitutes semantic representation in body-selective regions.

Although several fMRI and single-unit recording studies have measured responses in high-level visual regions to object pairs^49–52^ and proposed a computational framework to explain those responses^53–55^, these studies used only a handful of pairs of objects that were manually selected from many possible pairs. They thus remained silent about the extent to which each possible object pair drives neural responses. Our co-occurrence analysis examined the co-occurrence frequencies across as many as 66 pairs of object categories in natural scene images and extracted major co-occurrence patterns associated with the EBA response in a data-driven manner. As a result, we provide new evidence regarding the object category co-occurring with human bodies that potentially determines the magnitude of the EBA response and find that EBA encodes contextually defined information: human body motion speed implied in a static scene.

Previous fMRI studies offered evidence for the representation of implied motion in the brain. Some studies have shown that static images of humans and animals in dynamic action more strongly activate the MT and MST than by those without implied motion^33–35^. One study reported a stronger MT response, as higher motion speed is implied in static images of inanimate objects^56^. However, these studies did not explicitly investigate the effect of implied motion on body-selective regions. We computed vertex-wise correlations with ratings of implied motion speed throughout the visual cortex and demonstrated that EBA represents the speed of human and animal body motion implied in hundreds of natural scene images. A similar pattern was observed even when our analysis was restricted to body-selective vertices outside anatomically defined MT and MST. This aligns with previous findings that EBA, which is anatomically close to motion-selective areas, represents dynamic body actions in short video clips^57–60^. Importantly, our results extend these findings by showing that EBA also encodes the speed of implied motion in static images, which is consistent with an EEG study identifying EBA as a likely source of differential responses to static images with and without implied motion^61^. Notably, FBA— anatomically distinct from both the motion-selective areas and EBA—also showed strong correlations with the speed of implied motion. This highlights a novel aspect of FBA involvement in processing implied body motion. In view of these findings, the representation of implied motion is more widely distributed in the brain than previously reported and points to a broader functional role of EBA and FBA in representing human body motion inferred from accompanying objects and bodies themselves rather than merely responding to the presence of human bodies in natural scenes.

The notion that EBA and FBA encode implied human body motion is further supported by variance partitioning, which controls for other body-related features, such as body size and the number of people. For most subjects, implied motion uniquely and best explained the responses of the largest proportion of vertices in the EBA and FBA. Nevertheless, the number of people and body size depicted in the images also uniquely explained the responses in distinct subregions of these areas. This finding mirrors results from recent studies identifying two clusters within the EBA, each representing either individual bodies or bodies of multiple people^40,62^. However, these clusters encompassed almost the entire EBA. We argue that the apparent extent of the cluster reflecting the number of people may have been overestimated, as the contributions of multiple body-related features were not disentangled in these studies. Another finding from variance partitioning is that EBA and FBA differ in the second largest contributing body-related feature: the number of people for EBA and the body size for FBA. This regional difference is in line with previous studies showing that these two regions differ in several dimensions ^62–66^. We speculate that the difference observed in our study stems partly from distinct patterns of structural and functional connectivity between these regions: the EBA is strongly connected with the superior parietal lobe in the dorsal stream, whereas the FBA is connected with the fusiform gyrus and inferior temporal cortex in the ventral stream^67^. As the superior parietal lobe plays a role in shifting spatial attention^68,69^, which may be necessary to count the number of people distributed over a scene image, EBA is more likely to represent that information. On the other hand, estimating the size of bodies requires the shape processing of these bodies supported by regions in the ventral stream, thus resulting in a pronounced representation of body size in the FBA. Taken together, the semantic representations of EBA and FBA reported in recent studies^24^ are jointly shaped by several body-related features, where implied motion contributes most strongly in both regions, and the relative importance of these features differs across regions.

A significant gap still remains toward a comprehensive understanding of the semantic representations in EBA and FBA, as indicated by the fact that our linear regression model incorporating the three body-related features did not successfully explain all vertices. This is natural since the features we derived from the NSD images do not fully capture all aspects of human bodies in natural scenes. For example, emotion and social information expressed in the human body, which was previously reported to influence responses in the EBA and FBA^70–73^, may be a key factor that potentially improves the predictability of these regions. As it is inherently difficult to derive emotion-related features from NSD images that mostly contain emotionally neutral and nonsocial scenes, future studies should employ diverse sets of natural images and construct a model incorporating affective^74^, social^73,75^ and postural features associated with emotion categories^76,77^, as well as body-related features used in the present study to better interpret the semantic content represented in body-selective regions. Additionally, while most of our analyses were conducted at the single-vertex level, it is likely that distributed response patterns across multiple vertices encode body-related information, such as body posture, as shown in previous studies^72,78,79^. Further work is necessary to understand how information at different spatial scales is integrated to form semantic representations in the brain and achieve stable scene recognition.

Overall, we propose a co-occurrence analysis framework that moves one step further from the traditional view of category selectivity but still preserves the interpretability of neural representations in a target region. We believe that this framework provides substantial benefits to the field of cognitive neuroscience, as recent methodological advances have made it increasingly challenging to interpret results owing to the complexity of features used to model brain responses. We also show that this framework is indeed effective at understanding the contents of semantic representations in body-selective regions. Future studies could apply our framework to other category-selective regions and extend it to a broader range of categories to gain a deeper understanding of high-level semantic representations in the brain.

## Materials and methods

### Large-scale fMRI data

We utilized the Natural Scenes Dataset (NSD)^12^, which includes high-quality fMRI responses to a wide array of color natural scene images. These images are extracted from the Microsoft Common Objects in Context (COCO) database^28^. Eight NSD subjects observed 9000–10000 images in a 7T fMRI scanner, each of which was repeatedly presented up to three times throughout 40 scan sessions. Each image was 8.4° × 8.4° and was presented for 3 s. The subjects were asked to fixate on a centrally presented small dot and indicate by pressing a button whether they had seen each image before.

NSD provides whole-brain fMRI responses as beta values obtained from general linear modeling with optimized denoising and regularization methods (GLMsingle)^80^. We used beta values called fithrf_GLMdenoise_RR defined on the cortical native surface space of each subject, which were z-scored within each scan session. To restrict our analysis to a set of vertices with reliable signals, the noise ceiling signal-to-noise ratio (NCSNR) metric was used to subsample the vertices with NCSNR > 0.2, in accordance with a previous study^81^. After this subsampling process, we extracted response patterns for body-selective regions via the ROI definition and statistical contrast (t value) maps derived from functional localizer experiments in the NSD. Specifically, we defined the body-selective regions, extrastriate body area (EBA) and fusiform body area (FBA), as a group of vertices with t values > 1. We also defined motion-selective regions, i.e., the middle temporal (MT) and middle superior temporal (MST) areas, for each subject via anatomical definitions based on the HCP atlas (HCP_MMP1)^82^ projected onto the subject’s native space. Unless otherwise specified, our analyses were based on the mean response computed across repetitions of each image presentation.

### Co-occurrence analysis

To understand the semantic context that determines EBA responses to natural scene images, we constructed an encoding model using text descriptions of those images and their embeddings obtained from an LLM. Using this model, we predicted EBA responses to 73,000 NSD images and investigated how these predicted responses are associated with co-occurring word pairs in a set of text descriptions.

#### Encoding models based on text descriptions of natural scene images

We trained an encoding model that predicts the EBA response to a natural scene image given the corresponding caption provided by a human annotator (Figure 1B). Each NSD image has five captions, but we only used the first of these five captions included in the MS COCO database^27^. We then fed these captions into an LLM (all-mpnet (all-mpnet-base-v2)^30^ to extract 768-dimensional embeddings via the Sentence-Transformers library, in accordance with a previous study^24^. The embeddings served as predictors, allowing us to construct an encoding model using fractional ridge regression^83^ according to the following procedure: first, we held out as a test set the caption embeddings of 1000 images that were viewed by all eight subjects in the NSD experiment. The remaining embeddings (associated with images uniquely observed by each NSD subject) were subsequently used as a training set to estimate the weights and the best regularization hyperparameter of a ridge regression model for each NSD subject by setting regularization fraction values ranging from 0.1 to 1 in steps of 0.1. The trained encoding model was applied to the held-out embeddings to evaluate prediction accuracy by computing the Pearson correlation coefficient between the predicted and actual EBA responses averaged across repetitions for each image.

#### Defining a co-occurrence matrix from captions

A co-occurrence matrix typically consists of nonnegative integer values corresponding to the frequency of a word pair co-occurring in large-scale sentences. To define a co-occurrence matrix for the captions of the NSD images, we used multiple object categories at the superordinate level in the MS COCO dataset and their co-occurrence frequency in a group of captions. Specifically, we constructed a 12*12 matrix based on the co-occurrence frequency of 12 superordinate categories (accessory, animal, appliance, electronic, food, furniture, indoor, kitchen, outdoor, person, sports, and vehicle) derived from 91 basic COCO categories^28^, with two important modifications from the original category classification. First, we included several person-related words (man, woman, boy, girl, baby, child) as members of the “person” category. The reason for this modification is that these specific words are much more frequently used by human annotators to describe the presence of a person in images than the word “person”. Second, any words that constitute the four phrases included in the “sports” category (sports ball, baseball bat, baseball glove and tennis racket) are also considered members of the “sports” category (e.g., “baseball” and “bat” from “baseball bat”) because these constituent words are more frequently used in captions than are the phrases themselves. These modifications resulted in 12 superordinate categories derived from 100 basic categories (see Supplementary Table 1 for details).

Using these modified COCO object categories, we constructed multiple co-occurrence matrices from the captions associated with the NSD images. The construction process involves several steps. In the first step, we extracted the first of the five captions for each NSD image (as done in training encoding models) and fed the corresponding LLM (all-mpnet-base-v2) embeddings into the trained encoding models for each of the eight subjects, yielding predicted EBA responses to the 73,000 images for each subject (Figure 1C, left). In the second step, we sorted the 73,000 captions in descending order of the predicted EBA responses averaged across subjects and partitioned them into 73 groups of 1000 captions each, with the first group corresponding to the highest predicted EBA responses and the last to the lowest. The third step involves lemmatizing all nouns in the captions for each group using the WordNet lemmatizer in the nltk library, which converts inflected noun forms into their base form. This allowed us to accurately count all nouns of the 100 basic categories in the next step, regardless of their form in the captions. Finally, we counted the co-occurrence frequency of all 66 pairs of superordinate categories for the captions in each group, yielding 73 symmetric co-occurrence matrices that consisted of nonnegative integer values (Figure 1C, middle).

To extract major co-occurrence components closely linked to EBA responses in a data-driven manner, we performed Bayesian nonnegative matrix factorization on the data derived from the 73 co-occurrence matrices. Specifically, since the co-occurrence matrices are symmetric, we extracted their upper triangular portion, flattened it into a vector, and horizontally stacked the vectors to obtain a matrix of size 66*73 (Figure 1C, right). This matrix was decomposed into two different matrices: basis and coefficient matrices. Each column in the basis matrix represents a major co-occurrence pattern inherent in the original matrices, and each row in the coefficient matrix reflects the extent to which each co-occurrence pattern contributes to reconstructing the original matrix. By examining the 73 values in each row, we can assess how strongly each major co-occurrence pattern is associated with the 73 groups sorted based on EBA responses, thereby capturing its relationship with the magnitude of EBA responses. We performed this factorization while varying the number of components from 2 to 10 in increments of 1 and determined the optimal number of components as the one that minimizes the Bayesian information criterion (BIC) under the assumption of a Poisson distribution. The BIC was defined as follows:

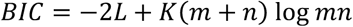

where m and n denote the number of rows and columns of the original matrix, respectively, and K indicates the number of components. The likelihood L was defined as follows:

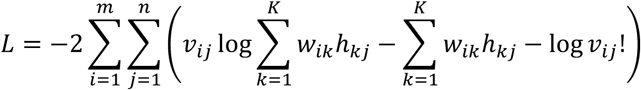

where v, w, and h are individual elements of the observed, basis, and coefficient matrices, respectively.

### Implied motion rating task

#### Participants

One of the authors and four naïve paid volunteers with corrected-to-normal vision participated in the behavioral experiment. Our experiments were conducted with permission from the Ethics Committee of the University of Tokyo. Written informed consent was obtained from all the participants prior to enrollment, and the Declaration of Helsinki guidelines were followed.

#### Image selection

Out of the 73000 natural scene images included in the NSD, we sampled a subset of those images for our rating experiment. We focused on 1000 shared images that were viewed by all eight subjects in the NSD experiment and used the image segmentation data (specifically the ‘person’ label) included in the original COCO category annotations to select images containing human bodies. We then subsampled the images for our rating experiment according to the following two selection criteria, which we believe facilitate the implied motion judgment of the person in an image. First, we ensured that all images do not contain animals or vehicles that potentially imply motion information unrelated to that of the person in an image. Second, we eliminated images in which the person appears too small in the background because inferring the implied motion of a person when the person occupies only a small portion of the image is difficult. After these two screening criteria were applied, 100 images were randomly selected from the resulting set. In addition to these 100 person images, we also sampled animal and vehicle images from the shared images labeled with the COCO superordinate categories ‘animal’ and ‘vehicle’. We again subsampled 100 images for each category based on the same criteria as above, ensuring that all the animal (vehicle) images did not contain people or vehicles (animals) and that the animals and vehicles in all the images did not appear too small. For the vehicle category, only 88 images met the criteria. As a result, a total of 288 natural images were selected to conduct an implied motion rating task for three categories.

#### Procedure

Our rating experiment consisted of three sessions in total, one for each target category (person, animal, vehicle), and the order of these sessions was counterbalanced across participants. Each session started with a passive-viewing session in which participants observed all images of a target category, each presented for 1 second, without response, so that they were aware of the dynamic range of motion information implied in the images^56^, making it easier for participants to rate the implied motion speed of a target category in subsequent rating blocks. In the next four blocks, participants were presented with an image for 3 seconds and evaluated the speed of implied motion of a target category for each image on a scale of 1 to 5. We provided participants with clear rating criteria: 1 indicates static motion, and 5 corresponds to the fastest motion they had previously seen in the passive-viewing block. We also instructed participants to give a rating within 3 secs of image presentation by considering the overall scene context in which the target category is present rather than relying solely on the semantic category of an image (e.g., we expected them to assign different ratings to an image of a surfer riding a wave and an image of a surfer just standing on the shore). To evaluate the within-participant reliability of ratings in a subsequent analysis, each image was presented twice throughout the four blocks; thus, the participants completed a total of 200 trials for the person and animal categories (50 trials per block) and 176 trials for the vehicle category (44 trials per block), with a few minutes of rest after two blocks. Prior to these main blocks, we also conducted a practice block in which participants rated the implied motion speed of 20 NSD images for each target category that were not included in the 288 images used in the main experiment based on the same criteria as above. Each image was presented on a monitor (BenQ XL2546X-B) with a size of 8.4°×8.4° at a viewing distance of 50 cm, followed by a 1-sec blank gray background, in accordance with the conditions of the NSD experiment^12^.

#### Rating reliability

We assessed two types of reliability for the ratings from our task: within-participant reliability, computed as Spearman’s rank correlation between ratings from the first and second image presentations for each participant, and between-participant reliability, computed as Spearman’s rank correlation between ratings averaged across two repetitions for each pair of participants. We confirmed high within-participant and between-participant reliability (Supplementary Figure 2). Thus, we averaged the ratings across repetitions and participants and used these averaged ratings for subsequent analyses.

### Correlation analysis using body-related image features

To test whether EBA responses to natural images are associated with the implied motion of a target category in those images, we first fitted a linear mixed-effects model that explains the EBA responses averaged across vertices using the implied motion ratings separately for each category (person, animal, vehicle). We included both random slopes and random intercepts for each subject, which capture individual differences across NSD subjects. The model is defined as:

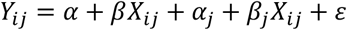

where *X_ij_* is the implied motion rating for image *i* (common across subjects *j*) and *Y_ij_* is the EBA response to image *i* for subject *j*. *α* and *β* represent the fixed-effect intercept and slope, respectively, while *α_j_* and *β_j_* denote the random intercept and slope for subject *j*. For each category, we assess the significance of the correlation between the EBA response and implied motion rating based on the p value of the fixed-effect slope.

To identify the cortical location representing the implied body motion, we additionally computed the correlation between the implied motion ratings and brain responses across vertices in the entire visual cortex. These vertices were defined via an ROI mask provided in the NSD (called the ‘stream’ covering the early, intermediate, and high-level visual cortex in the ventral and dorsal streams), allowing us to test whether the observed correlation profile is specific to body-selective regions. Spearman’s rank correlations between the vertex-wise responses and the implied body motion ratings for each image were computed for each NSD subject. To evaluate the statistical significance of the correlation for each vertex, we performed a one-sided permutation test with 1000 randomizations. For each randomization, we permuted the stimulus labels and computed a correlation, yielding a null distribution of correlation coefficients. P-values were defined as the proportion of correlations exceeding the observed correlation coefficient and were corrected for multiple comparisons using the Benjamin`‒`Hochberg false discovery rate (FDR) procedure (*p* < 0.05).

Furthermore, we computed additional body-related and low-level features (body size, number of people, distance from the center, and RMS contrast; see below for the definition and the rationale behind this feature selection) for each image to determine whether these features unrelated to implied motion could also account for the response in the body-selective regions. We again computed the correlation between these additional features and brain responses in the body-selective regions by using Spearman’s rank correlation for the number of people and the Pearson correlation for the other features.

The same statistical procedure as above was employed, but two-sided tests were used in this analysis because, for distance from the center, we expected a negative correlation with brain response (i.e., higher response associated with smaller distance from the image center). The correlation matrix for all pairs of features is provided in Supplementary Figure 6.

#### Body size

Human bodies covering a large amount of space in an image possibly elicit greater responses in body-selective regions^4^. To compute the body size for each image, we used a text-based segmentation model called ‘Grounded Segment Anything’^42^. This model leverages the strengths of two machine learning models: grounding DINO^84^, which identifies the presence of an object specified by text input, and Segment Anything^85^, which performs segmentation of objects and people within an image. For each of the 100 images used in the rating experiment, we segmented people by providing the text input “person” to the model and computed the human body size, which was defined as the number of pixels corresponding to the human body in each image. If multiple people were present in an image, we computed the total body size by summing the body sizes of all individuals.

#### Number of people

One recent study revealed that EBA has two distinct clusters, with one cluster representing a group of people and the other representing a single person in NSD images^40^, suggesting the possibility that the number of people present in an image determines EBA responses. To test this possibility, we extracted the segmentation labels provided by the Grounded Segment Anything for each of the 100 images and counted the number of “person” labels.

#### Distance from the center

Given that different parts of the body-selective regions exhibit distinct eccentricity biases^36^, the spatial distance between the human body and the center of an image (where a fixation point was in the NSD experiment) could influence the magnitude of EBA responses. To explore this, we again used the output from Grounded Segment Anything, particularly the bounding box of the largest human body in each image. The distance was computed as the Euclidean distance between the center of an image and the center of the bounding box.

#### RMS contrast

A qualitative assessment of images potentially implying faster motion (e.g., an image of a skier or an airplane against a uniform blue sky) raised the possibility that EBA responses can be modulated by spatial frequency information, rather than the implied motion of an image. We thus converted the 100 images to grayscale values and computed the root mean square (RMS) contrast of the pixel intensities for each image as a proxy for the spatial frequency. This value tends to be lower when large uniform regions occupy the image.

### Variance partitioning

To examine the unique and shared contributions of three major body-related features (implied motion, body size, and number of people) to the prediction of brain responses for each vertex, we performed variance partitioning on the basis of seven linear regression models. Specifically, we quantified the amount of variance uniquely explained by each feature, as well as the variance jointly explained by two or all three features. Explained variance was quantified using the coefficient of determination (R^2^), which was computed for each regression model via 5-fold cross-validation. For each fold, the linear regression model was trained on 80% of the data and tested on the remaining 20%. The cross-validated R^2^ values obtained from the test sets were averaged across the five folds to robustly estimate the model performance. On the basis of the seven cross-validated R^2^ values 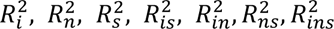 for implied motion, number of people, body size, a pair of implied motion and body size, a pair of implied motion and number of people, a pair of body size and number of people, and a combination of all three features, respectively), we computed the unique variance for each feature and the shared variance for all possible combinations of features. For instance, the uniquely explained variance for implied motion (*U_i_*) was computed as follows:

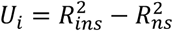

The variance shared by the implied motion and number of people (*S_in_*) and among all three features (*S_ins_*), was computed via the following equations:

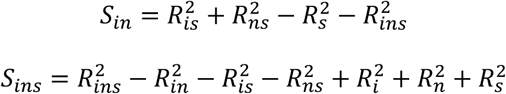

Statistical significance was assessed using a permutation test as described above. We used the same randomization for all seven regression models and evaluated the significance of the observed cross-validated R^2^ values on the basis of the significance threshold (*p* < 0.05; FDR-corrected) defined by a null distribution of R^2^ values for each vertex. We then extracted the vertices for which at least one of the seven regression models achieved significant prediction accuracy and computed the proportion of vertices that were best explained by each feature or combination of multiple features.

### Data availability

The Natural Scene Dataset is publicly available at http://naturalscenesdataset.org. The data of the behavioral experiment will be made available via a GitHub repository upon publication of the manuscript.

### Code availability

The code for the analyses reported here will be made available via a GitHub repository upon publication of the manuscript.

## Acknowledgments

This work was supported by JSPS KAKENHI Grant Number 23KJ0477 (R.Y.). Collection of the NSD dataset was supported by NSF IIS-1822683 and NSF IIS-1822929.

## Author contributions

R.Y., M.S., A.Y., and K.A. conceived the study and designed the behavioral experiment. R.Y. collected the behavioral data, wrote the analysis scripts, performed all analyses, and drafted the manuscript. K.A., M.S., and A.Y. supervised the study. R.Y., M.S., A.Y., and K.A. reviewed and approved the final manuscript.

## Competing interests

The authors declared no competing interests.

## Supplementary Information

**Supplementary Table 1.** List of 12 superordinate categories defined in the MS COCO dataset and 100 subordinate categories used in the co-occurrence analysis.

| *accessory* | *animal* | *appliance* | *electronic* | *food* | *furniture* |
| --- | --- | --- | --- | --- | --- |
| backpack | bear | blender | cell phone | apple | bed |
| eye glasses | bird | microwave | keyboard | banana | chair |
| handbag | cat | oven | laptop | broccoli | couch |
| hat | cow | refrigerator | mouse | cake | desk |
| shoe | dog | sink | remote | carrot | dining table |
| suitcase | elephant | toaster | tv | chair | door |
| tie | giraffe |  |  | donut | mirror |
| umbrella | horse |  |  | hot dog | potted plant |
|  | sheep |  |  | orange | toilet |
|  | zebra |  |  | pizza | window |

| *indoor* | *kitchen* | *outdoor* | *person* | *sports* | *vehicle* |
| --- | --- | --- | --- | --- | --- |
| book | bottle | bench | baby | baseball | airplane |
| clock | bowl | fire hydrant | boy | bat | bicycle |
| hair brush | cup | parking meter | child | ball | boat |
| hair drier | fork | stop sign | girl | frisbee | bus |
| scissors | knife | street sign | man | glove | car |
| teddy bear | plate | traffic light | person | kite | motorcycle |
| toothbrush | spoon |  | woman | racket | train |
| vase | wine glass |  |  | skateboard | truck |
|  |  |  |  | skis |  |
|  |  |  |  | snowboard |  |
|  |  |  |  | sports |  |
|  |  |  |  | surfboard |  |
|  |  |  |  | tennis |  |

**Supplementary Table 2.**
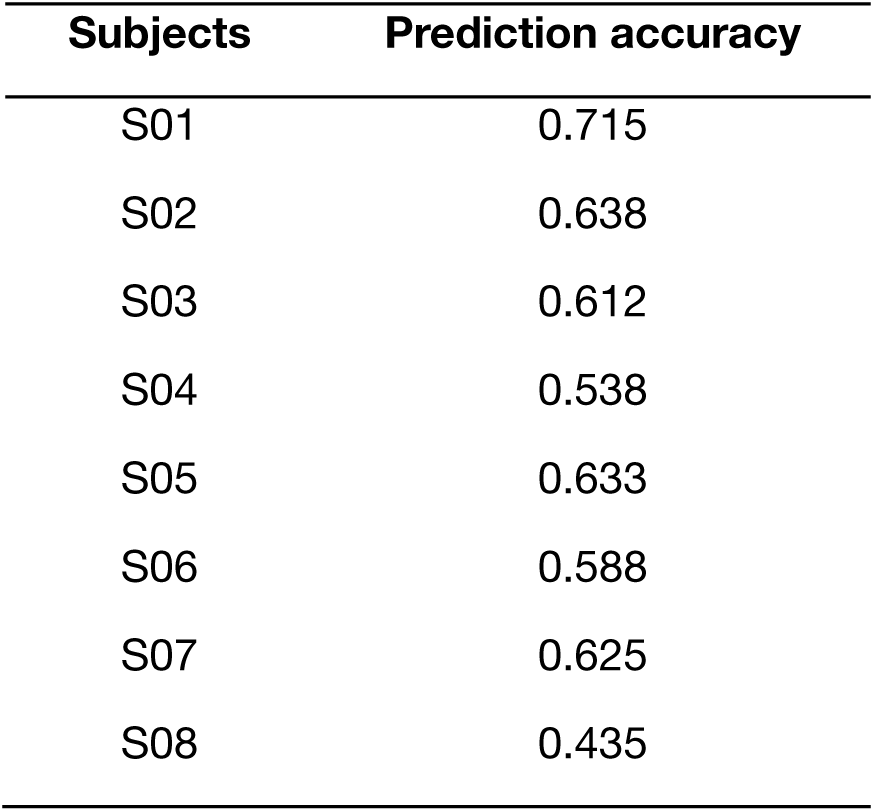
Prediction accuracy of the caption-based encoding model for each NSD subject defined as Pearson correlation coefficient between observed and predicted mean EBA responses.

**Supplementary Figure 1.**
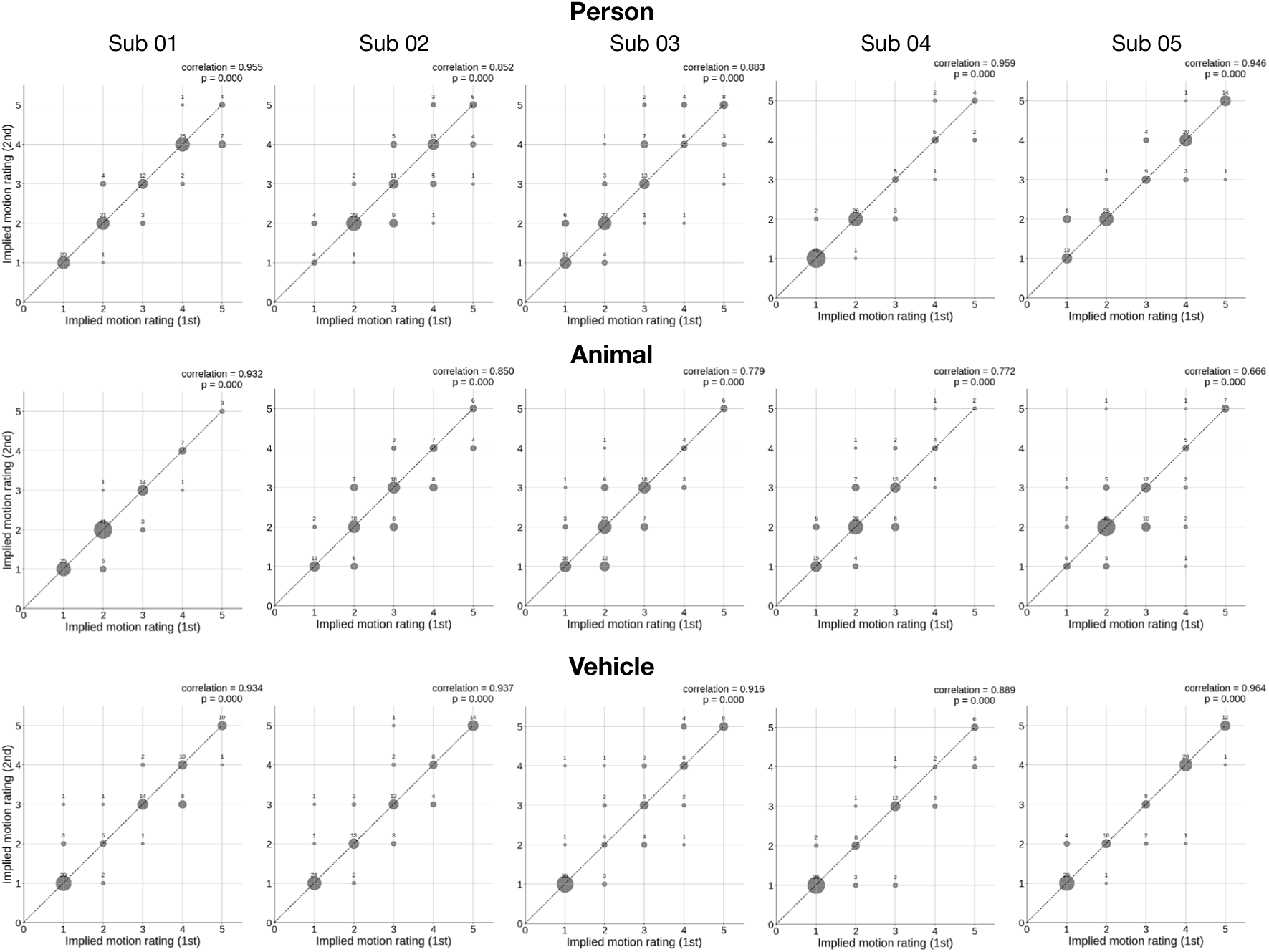
Within-participant reliability (Spearman rank correlation of responses between the first and second image presentations) in the rating experiment for each category. Each bubble plot shows the frequency of ratings across two repetitions, with larger bubbles reflecting more frequent responses.

**Supplementary Figure 2.**
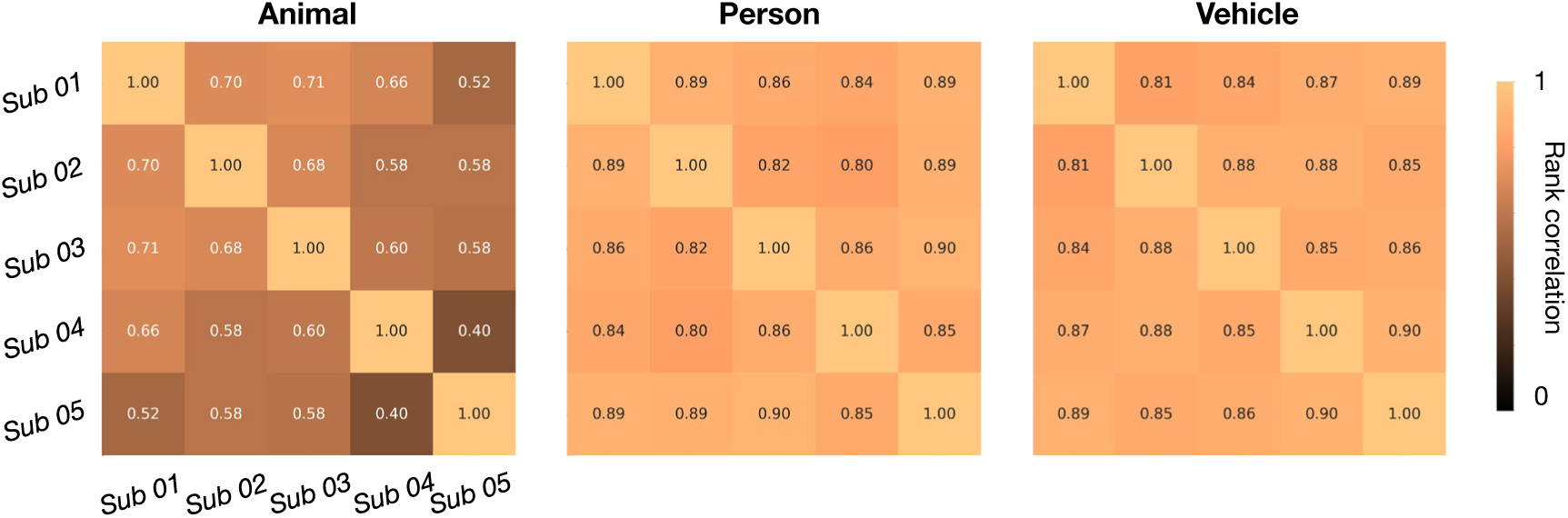
Between-participant reliability (Spearman rank correlations of responses between all pairs of five subjects) in the rating experiment for each category.

**Supplementary Figure 3.**
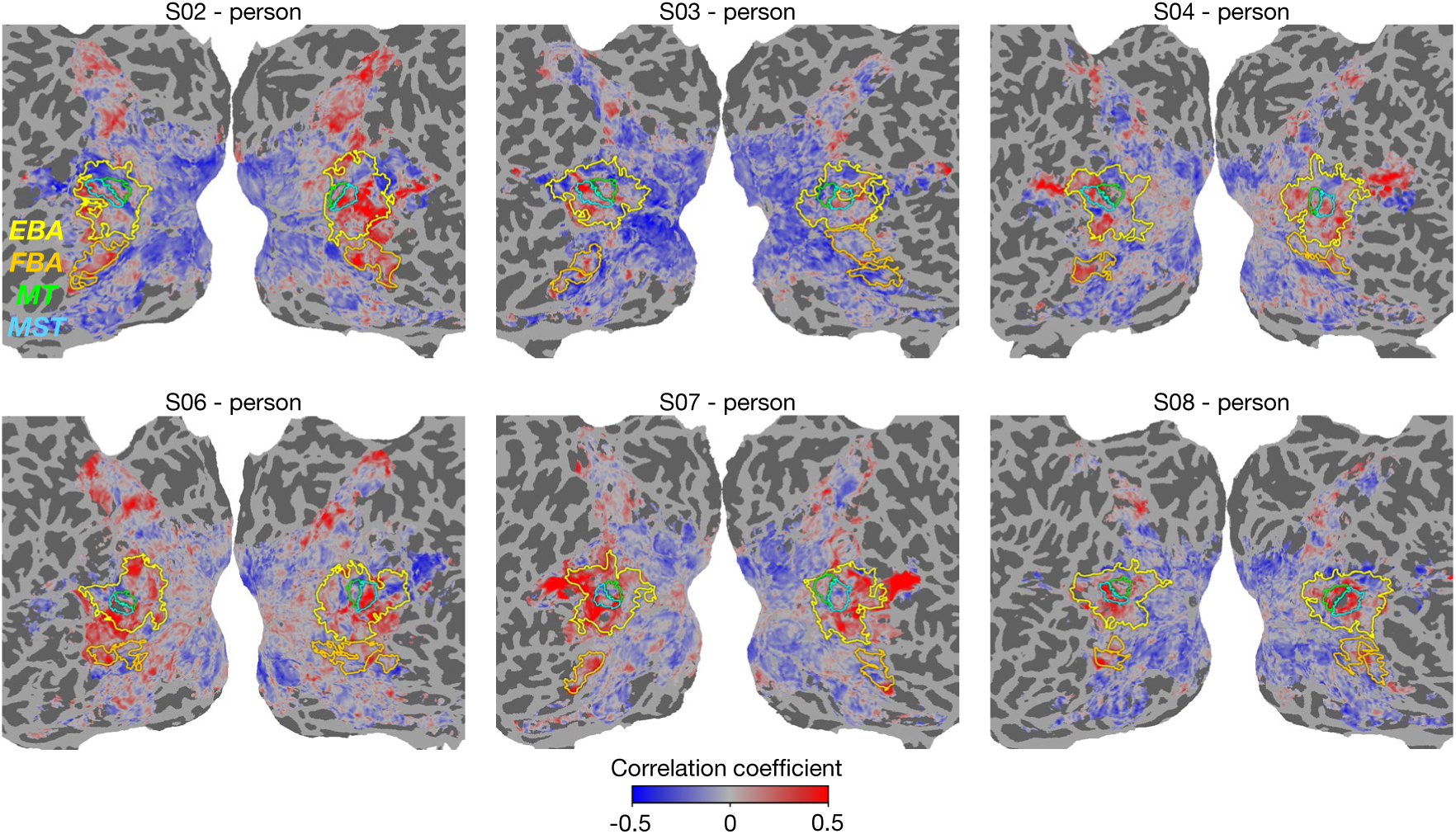
Cortical surface maps of the correlation between vertex-wise responses and implied motion ratings of person images for the remaining 6 NSD subjects. The maps for subject 01 and 05 are shown in Figure 4A.

**Supplementary Figure 4.**
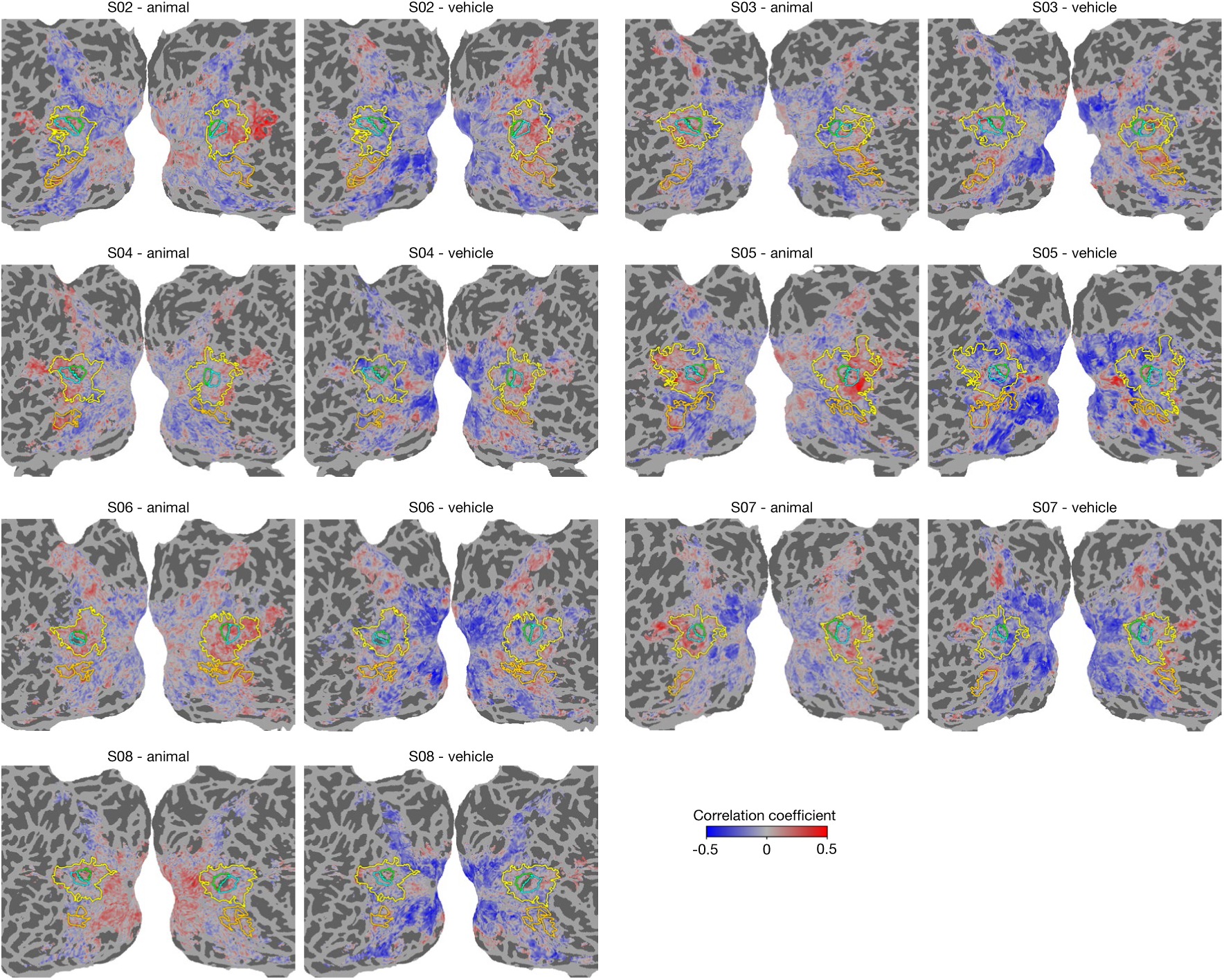
Cortical surface maps of the correlation between vertex-wise responses and implied motion ratings of animal and vehicle images for the remaining 7 NSD subjects. The maps for subject 01 are shown in Figure 4B.

**Supplementary Figure 5.**
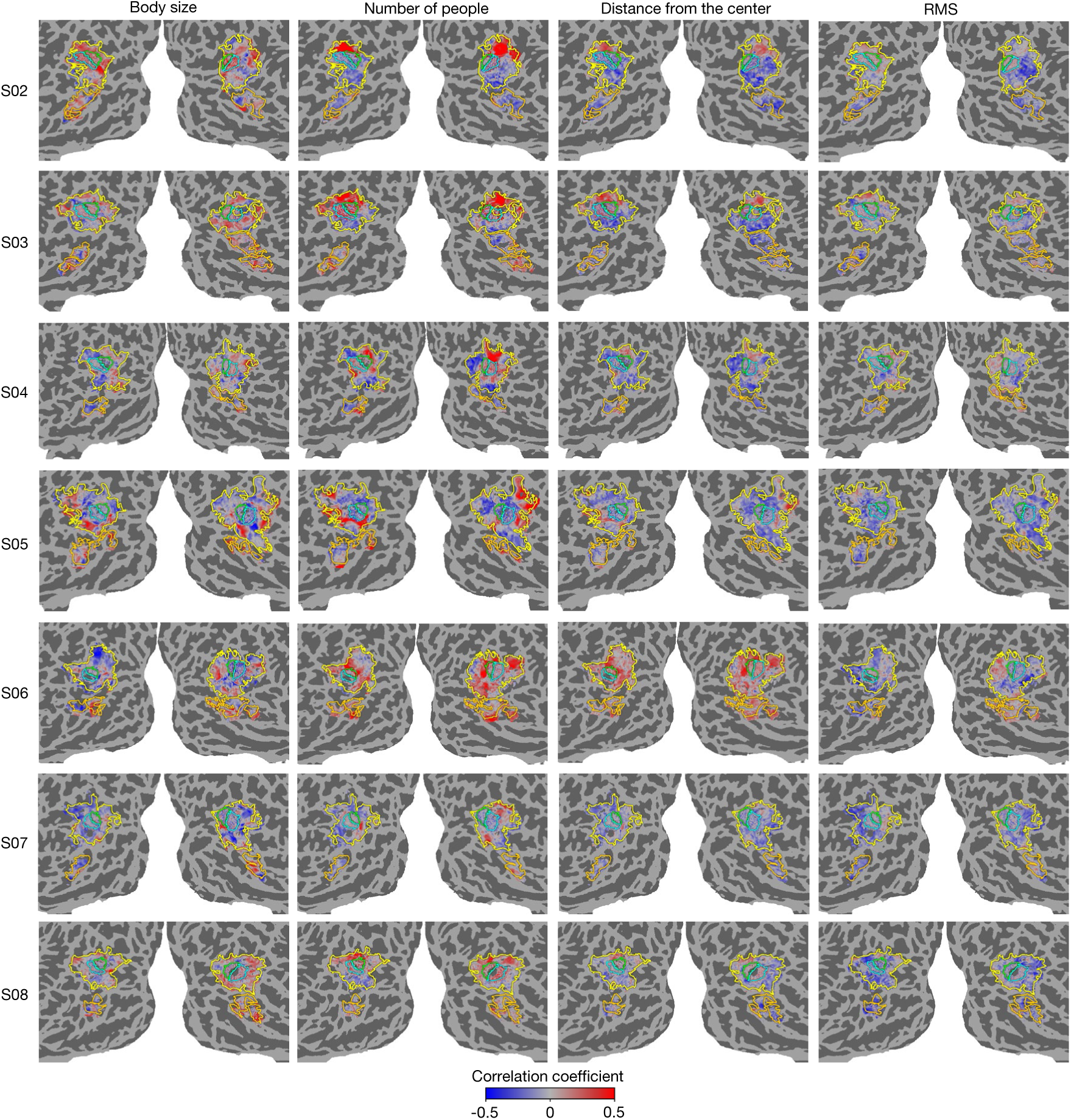
Flattened cortical surface maps of correlation coefficients with the three body-related features (body size, number of people, and distance from the center) and one low-level visual feature (RMS contrast) for the remaining 7 NSD subjects. The maps for subject 01 are shown in Figure 4B.

**Supplementary Figure 6.**
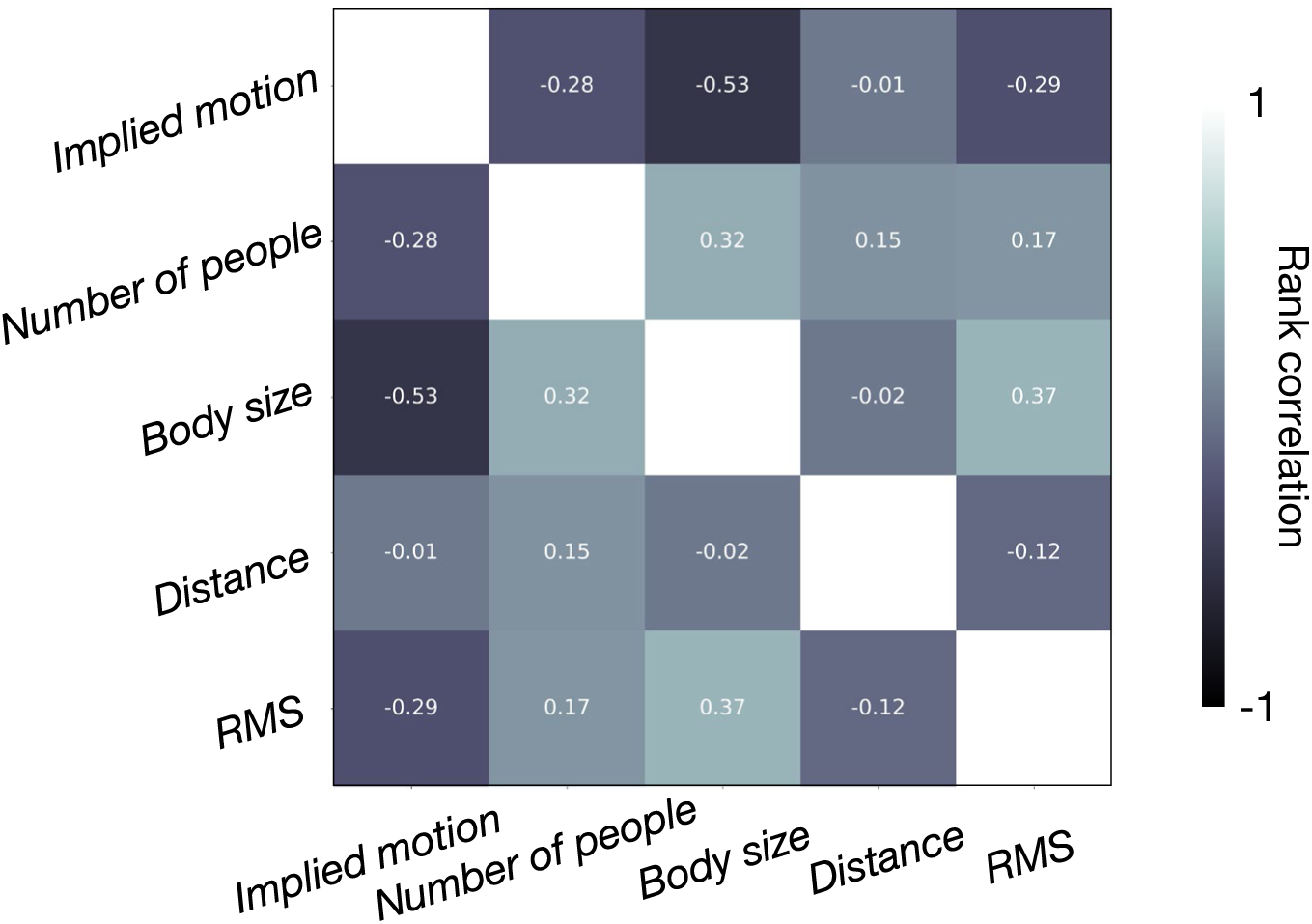
Rank correlation between all pairs of features.

**Supplementary Figure 7.**
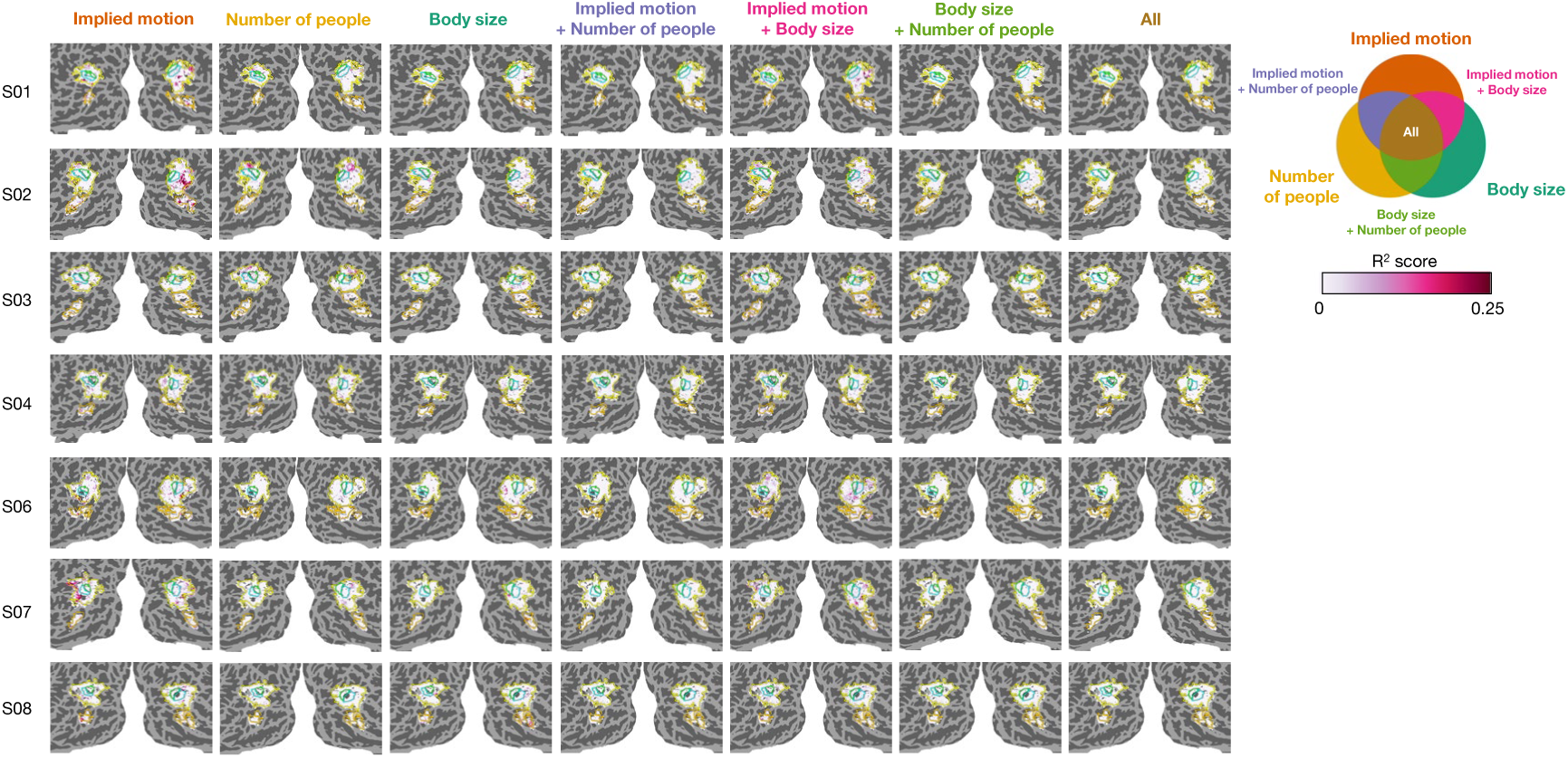
Cortical surface maps of variance uniquely explained by each individual feature and variance jointly explained by a pair or triplet of features for the remaining 7 NSD subjects. The maps for subject 05 are shown in Figure 6.

**Supplementary Figure 8.**
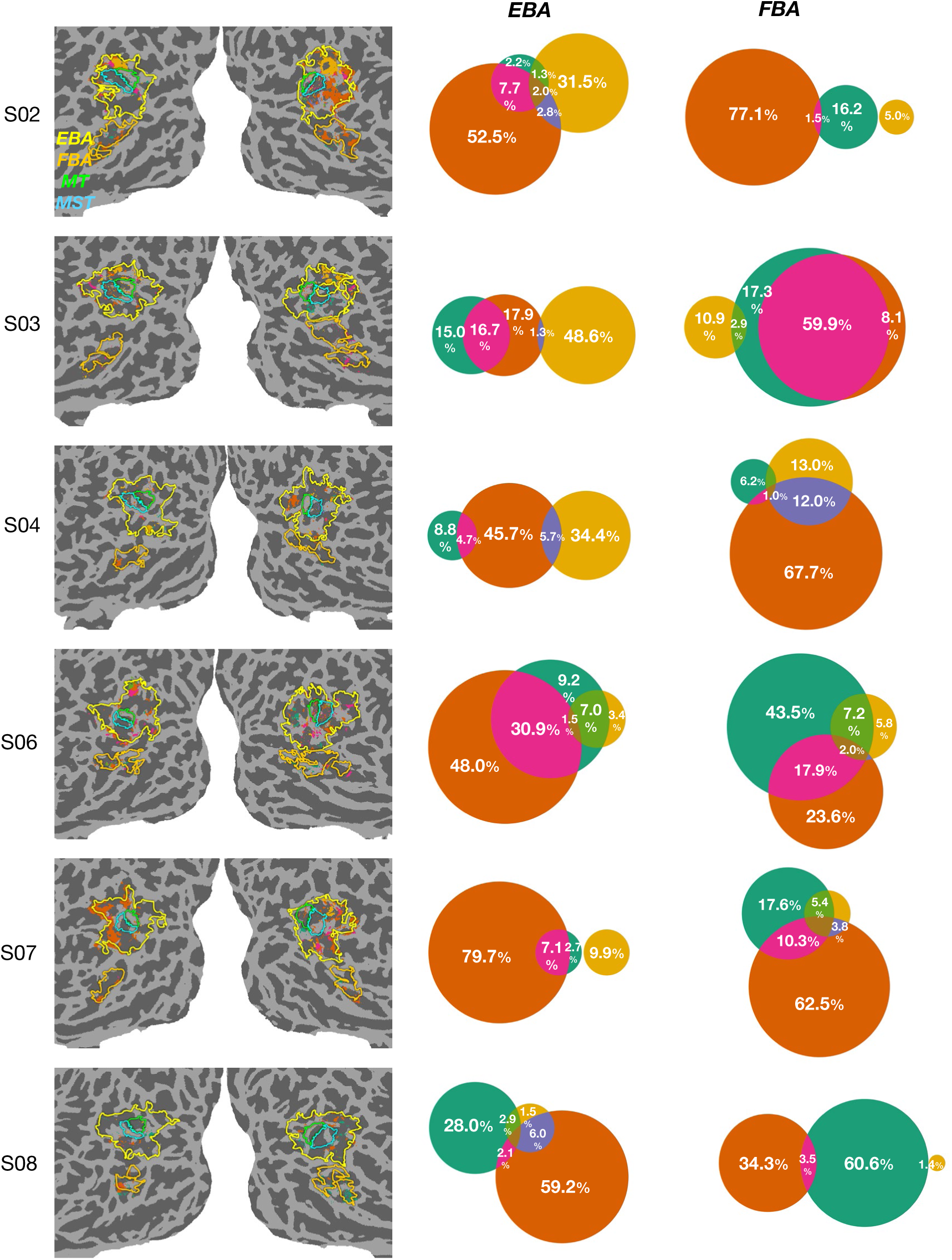
Cortical surface maps of the largest variance partition for the remaining 6 NSD subjects. The Venn diagrams show the proportion of significantly predicted vertices best explained by each feature (note that the size of each area is not scaled to reflect exact proportions).

## Notes

### Competing Interest Statement

The authors have declared no competing interest.

## References

1. Peelen, M. V. & Downing, P. E. The neural basis of visual body perception. Nat. Rev. Neurosci. 8, 636–648 (2007).

2. Downing, P. E. & Peelen, M. V. The role of occipitotemporal body-selective regions in person perception. Cogn. Neurosci. 2, 186–203 (2011).

3. Downing, P. E., Jiang, Y., Shuman, M. & Kanwisher, N. A cortical area selective for visual processing of the human body. Science 293, 2470–2473 (2001).

4. Taylor, J. C., Wiggett, A. J. & Downing, P. E. Functional MRI analysis of body and body part representations in the extrastriate and fusiform body areas. J. Neurophysiol. 98, 1626–1633 (2007).

5. Urgesi, C., Berlucchi, G. & Aglioti, S. M. Magnetic stimulation of extrastriate body area impairs visual processing of nonfacial body parts. Curr. Biol. 14, 2130–2134 (2004).

6. Peelen, M. V., Wiggett, A. J. & Downing, P. E. Patterns of fMRI activity dissociate overlapping functional brain areas that respond to biological motion. Neuron 49, 815–822 (2006).

7. Michels, L., Lappe, M. & Vaina, L. Visual areas involved in the perception of human movement from dynamic form analysis. Neuroreport 16, 1037–1041 (2005).

8. Vangeneugden, J., Peelen, M. V., Tadin, D. & Battelli, L. Distinct neural mechanisms for body form and body motion discriminations. J. Neurosci. 34, 574–585 (2014).

9. Kayser, C., Körding, K. P. & König, P. Processing of complex stimuli and natural scenes in the visual cortex. Curr. Opin. Neurobiol. 14, 468–473 (2004).

10. Nastase, S. A., Goldstein, A. & Hasson, U. Keep it real: rethinking the primacy of experimental control in cognitive neuroscience. Neuroimage 222, 117254 (2020).

11. Hebart, M. N. et al. THINGS-data, a multimodal collection of large-scale datasets for investigating object representations in human brain and behavior. Elife 12, (2023).

12. Allen, E. J. et al. A massive 7T fMRI dataset to bridge cognitive neuroscience and artificial intelligence. Nat. Neurosci. 25, 116–126 (2021).

13. Lahner, B. et al. Modeling short visual events through the BOLD moments video fMRI dataset and metadata. Nat. Commun. 15, 6241 (2024).

14. Krizhevsky, A., Sutskever, I. & Hinton, G. E. ImageNet classification with deep convolutional neural networks. Commun. ACM 60, 84–90 (2017).

15. Vaswani, A., et al. Attention Is All You Need. *arXiv [cs.CL]* (2017).

16. Dosovitskiy, A., et al. An image is worth 16×16 words: Transformers for image recognition at scale. *arXiv [cs.CV]* (2020).

17. Bashivan, P., Kar, K. & DiCarlo, J. J. Neural population control via deep image synthesis. Science 364, (2019).

18. Khosla, M., Ratan Murty, N. A. & Kanwisher, N. A highly selective response to food in human visual cortex revealed by hypothesis-free voxel decomposition. Curr. Biol. 32, 4159–4171.e9 (2022).

19. Jain, N. et al. Selectivity for food in human ventral visual cortex. Commun Biol 6, 175 (2023).

20. Contier, O., Baker, C. I. & Hebart, M. N. Distributed representations of behaviour-derived object dimensions in the human visual system. Nat. Hum. Behav. 1–15 (2024).

21. Takagi, Y. & Nishimoto, S. High-resolution image reconstruction with latent diffusion models from human brain activity. *bioRxiv* 2022.11.18.517004 (2023) doi:10.1101/2022.11.18.517004.

22. Horikawa, T. Mind captioning: Evolving descriptive text of mental content from human brain activity. *bioRxiv* 2024.04.23.590673 (2024) doi:10.1101/2024.04.23.590673.

23. Richard, A., et al. A generative framework to bridge data-driven models and scientific theories in language neuroscience. *arXiv [cs.CL]* (2024).

24. Doerig, A. et al. High-level visual representations in the human brain are aligned with large language models. Nat. Mach. Intell. 1–15 (2025).

25. Pennington, J., Socher, R. & Manning, C. D. GloVe: Global Vectors for Word Representation. in Proceedings of the 2014 Conference on Empirical Methods in Natural Language Processing (EMNLP) 1532–1543 (2014).

26. Bojanowski, P., Grave, E., Joulin, A. & Mikolov, T. Enriching word vectors with subword information. *arXiv [cs.CL]* (2016).

27. Chen, X. et al. Microsoft COCO captions: Data collection and evaluation server. arXiv [cs.CV] (2015).

28. Lin, T.-Y. et al. Microsoft COCO: Common Objects in Context. *arXiv [cs.CV]* (2014).

29. Peelen, M. V. & Downing, P. E. Selectivity for the human body in the fusiform gyrus. J. Neurophysiol. 93, 603–608 (2005).

30. Song, K., Tan, X., Qin, T., Lu, J. & Liu, T.-Y. MPNet: Masked and Permuted Pre-training for Language Understanding. *arXiv [cs.CL]* (2020).

31. Freyd, J. J. The mental representation of movement when static stimuli are viewed. Percept. Psychophys. 33, 575–581 (1983).

32. Kourtzi, Z. But still, it moves. Trends Cogn. Sci. 8, 47–49 (2004).

33. Kourtzi, Z. & Kanwisher, N. Activation in human MT/MST by static images with implied motion. J. Cogn. Neurosci. 12, 48–55 (2000).

34. Senior, C. et al. The functional neuroanatomy of implicit-motion perception or representational momentum. Curr. Biol. 10, 16–22 (2000).

35. Senior, C., Ward, J. & David, A. S. Representational momentum and the brain: An investigation into the functional necessity of V5/MT. Vis. Cogn. 9, 81–92 (2002).

36. Weiner, K. S. & Grill-Spector, K. Not one extrastriate body area: using anatomical landmarks, hMT+, and visual field maps to parcellate limb-selective activations in human lateral occipitotemporal cortex. Neuroimage 56, 2183–2199 (2011).

37. Ferri, S., Kolster, H., Jastorff, J. & Orban, G. A. The overlap of the EBA and the MT/V5 cluster. Neuroimage 66, 412–425 (2013).

38. Kanwisher, N., McDermott, J. & Chun, M. M. The fusiform face area: a module in human extrastriate cortex specialized for face perception. J. Neurosci. 17, 4302–4311 (1997).

39. Epstein, R., Harris, A., Stanley, D. & Kanwisher, N. The parahippocampal place area: recognition, navigation, or encoding? Neuron 23, 115–125 (1999).

40. Luo, A., Henderson, M. M., Tarr, M. J. & Wehbe, L. BrainSCUBA: Fine-Grained Natural Language Captions of Visual Cortex Selectivity. in The Twelfth International Conference on Learning Representations (2023).

41. Schwarzlose, R. F., Swisher, J. D., Dang, S. & Kanwisher, N. The distribution of category and location information across object-selective regions in human visual cortex. Proc. Natl. Acad. Sci. U. S. A. 105, 4447–4452 (2008).

42. Ren, T., et al. Grounded SAM: Assembling open-world models for diverse visual tasks. *arXiv [cs.CV]* (2024).

43. Lescroart, M. D., Stansbury, D. E. & Gallant, J. L. Fourier power, subjective distance, and object categories all provide plausible models of BOLD responses in scene-selective visual areas. Front. Comput. Neurosci. 9, 135 (2015).

44. de Heer, W. A., Huth, A. G., Griffiths, T. L., Gallant, J. L. & Theunissen, F. E. The hierarchical cortical organization of human speech processing. J. Neurosci. 37, 6539– 6557 (2017).

45. Op de Beeck, H. P., Brants, M., Baeck, A. & Wagemans, J. Distributed subordinate specificity for bodies, faces, and buildings in human ventral visual cortex. Neuroimage 49, 3414–3425 (2010).

46. Bracci, S., Ietswaart, M., Peelen, M. V. & Cavina-Pratesi, C. Dissociable neural responses to hands and non-hand body parts in human left extrastriate visual cortex. J. Neurophysiol. 103, 3389–3397 (2010).

47. Bracci, S., Caramazza, A. & Peelen, M. V. Representational similarity of body parts in human occipitotemporal cortex. J. Neurosci. 35, 12977–12985 (2015).

48. Saxe, R., Jamal, N. & Powell, L. My body or yours? The effect of visual perspective on cortical body representations. Cereb. Cortex 16, 178–182 (2006).

49. Zoccolan, D., Cox, D. D. & DiCarlo, J. J. Multiple object response normalization in monkey inferotemporal cortex. J. Neurosci. 25, 8150–8164 (2005).

50. Cox, D., Meyers, E. & Sinha, P. Contextually evoked object-specific responses in human visual cortex. Science 304, 115–117 (2004).

51. Harel, A., Kravitz, D. J. & Baker, C. I. Deconstructing visual scenes in cortex: gradients of object and spatial layout information. Cereb. Cortex 23, 947–957 (2013).

52. Kim, J. G. & Biederman, I. Where do objects become scenes? Cereb. Cortex 21, 1738– 1746 (2011).

53. Bao, P. & Tsao, D. Y. Representation of multiple objects in macaque category-selective areas. Nat. Commun. 9, 1774 (2018).

54. Kliger, L. & Yovel, G. The functional organization of High-Level visual cortex determines the representation of complex visual stimuli. J. Neurosci. 40, 7545–7558 (2020).

55. Doostani, N., Hossein-Zadeh, G.-A. & Vaziri-Pashkam, M. The normalization model predicts responses in the human visual cortex during object-based attention. Elife 12, (2023).

56. Lu, Z., Li, X. & Meng, M. Encodings of implied motion for animate and inanimate object categories in the two visual pathways. Neuroimage 125, 668–680 (2016).

57. Downing, P. E., Peelen, M. V., Wiggett, A. J. & Tew, B. D. The role of the extrastriate body area in action perception. Soc. Neurosci. 1, 52–62 (2006).

58. Kable, J. W. & Chatterjee, A. Specificity of action representations in the lateral occipitotemporal cortex. J. Cogn. Neurosci. 18, 1498–1517 (2006).

59. Wiggett, A. J. & Downing, P. E. Representation of action in occipito-temporal cortex. J. Cogn. Neurosci. 23, 1765–1780 (2011).

60. Takahashi, H. et al. Enhanced activation in the extrastriate body area by goal-directed actions. Psychiatry Clin. Neurosci. 62, 214–219 (2008).

61. Proverbio, A. M., Riva, F. & Zani, A. Observation of static pictures of dynamic actions enhances the activity of movement-related brain areas. PLoS One 4, e5389 (2009).

62. Abassi, E. & Papeo, L. The representation of two-body shapes in the human visual cortex. J. Neurosci. 40, 852–863 (2020).

63. Foster, C. et al. Decoding subcategories of human bodies from both body- and face-responsive cortical regions. Neuroimage 202, 116085 (2019).

64. van de Riet, W. A. C., Grezes, J. & de Gelder, B. Specific and common brain regions involved in the perception of faces and bodies and the representation of their emotional expressions. Soc. Neurosci. 4, 101–120 (2009).

65. Ren, J. et al. Features and extra-striate body area representations of diagnostic body parts in anger and fear perception. Brain Sci. 12, 466 (2022).

66. Hodzic, A., Kaas, A., Muckli, L., Stirn, A. & Singer, W. Distinct cortical networks for the detection and identification of human body. Neuroimage 45, 1264–1271 (2009).

67. Zimmermann, M., Mars, R. B., de Lange, F. P., Toni, I. & Verhagen, L. Is the extrastriate body area part of the dorsal visuomotor stream? Brain Struct. Funct. 223, 31–46 (2018).

68. Vandenberghe, R., Gitelman, D. R., Parrish, T. B. & Mesulam, M. M. Functional specificity of superior parietal mediation of spatial shifting. Neuroimage 14, 661–673 (2001).

69. Corbetta, M., Shulman, G. L., Miezin, F. M. & Petersen, S. E. Superior parietal cortex activation during spatial attention shifts and visual feature conjunction. Science 270, 802– 805 (1995).

70. Marrazzo, G., Vaessen, M. J. & de Gelder, B. Decoding the difference between explicit and implicit body expression representation in high level visual, prefrontal and inferior parietal cortex. Neuroimage 243, p (2021).

71. Marsh, A. A. et al. The neural substrates of action identification. Soc. Cogn. Affect. Neurosci. 5, 392–403 (2010).

72. Poyo Solanas, M., Vaessen, M. & de Gelder, B. Computation-based feature representation of body expressions in the human brain. Cereb. Cortex 30, 6376–6390 (2020).

73. McMahon, E., Bonner, M. F. & Isik, L. Hierarchical organization of social action features along the lateral visual pathway. Curr. Biol. 33, 5035–5047.e8 (2023).

74. Abdel-Ghaffar, S. A. et al. Occipital-temporal cortical tuning to semantic and affective features of natural images predicts associated behavioral responses. Nat. Commun. 15, 5531 (2024).

75. Gandolfo, M. et al. Converging evidence that left extrastriate body area supports visual sensitivity to social interactions. Curr. Biol. 34, 343–351.e5 (2024).

76. Poyo Solanas, M., Vaessen, M. J. & de Gelder, B. The role of computational and subjective features in emotional body expressions. Sci. Rep. 10, 6202 (2020).

77. de Gelder, B. & Poyo Solanas, M. A computational neuroethology perspective on body and expression perception. Trends Cogn. Sci. 25, 744–756 (2021).

78. Marrazzo, G., De Martino, F., Lage-Castellanos, A., Vaessen, M. J. & de Gelder, B. Voxelwise encoding models of body stimuli reveal a representational gradient from low-level visual features to postural features in occipitotemporal cortex. Neuroimage 277, 120240 (2023).

79. Zhu, H. et al. Natural scenes reveal diverse representations of 2D and 3D body pose in the human brain. Proc. Natl. Acad. Sci. U. S. A. 121, e2317707121 (2024).

80. Prince, J. S. et al. Improving the accuracy of single-trial fMRI response estimates using GLMsingle. Elife 11, (2022).

81. Conwell, C., Prince, J. S., Kay, K. N., Alvarez, G. A. & Konkle, T. A large-scale examination of inductive biases shaping high-level visual representation in brains and machines. Nat. Commun. 15, 9383 (2024).

82. Glasser, M. F. et al. A multi-modal parcellation of human cerebral cortex. Nature 536, 171– 178 (2016).

83. Rokem, A. & Kay, K. Fractional ridge regression: a fast, interpretable reparameterization of ridge regression. Gigascience 9, giaa133 (2020).

84. Liu, S., et al. Grounding DINO: Marrying DINO with grounded pre-training for open-set object detection. *arXiv [cs.CV]* (2023).

85. Kirillov, A., et al. Segment Anything. *arXiv [cs.CV]* (2023).

